# GATA3-extension mutants rewire lineage identity in luminal breast cancer

**DOI:** 10.64898/2026.08.22.746471

**Authors:** Nivedhya Venas, Mrunal Ratnaparkhi, Kartik Majila, Stacey Jamieson, Igor Chernukhin, Shruthi Viswanath, Jason S. Carroll, Dimple Notani, Radhakrishnan Sabarinathan

**Affiliations:** National Centre for Biological Sciences, Tata Institute of Fundamental Research, Bengaluru, 560065, India; Cancer Research UK Cambridge Institute, Robinson Way, Cambridge, CB2 0RE, United Kingdom

## Abstract

GATA3, one of the most frequently mutated transcription factors in breast cancers, is a master regulator of luminal epithelial identity. Unlike many truncating and splice-site GATA3 mutations that disrupt one of the two zinc finger DNA-binding domains, C-terminal extension mutations (eGATA3) retain both of them. This raises the question of whether the eGATA3 mutants can alter transcriptional regulation and thereby exert distinct functional effects. Here, we integrate patient tumor-derived transcription and chromatin accessibility profiling with cell line derived genome-wide mapping of GATA3 occupancy and single-cell multiomics to define the regulatory consequences of eGATA3. We show that eGATA3 is associated with poor clinical outcome and remodels the luminal transcriptional program, leading to attenuation of estrogen-responsive transcription and progressive loss of luminal differentiation. Mechanistically, eGATA3 maintains widespread chromatin occupancy but redistributes GATA3 binding across different classes of regulatory elements, with preferential loss at AP-1 motif-enriched sites and gain at FOX-associated enhancers, along with coordinated remodeling of chromatin accessibility. These changes rewire the luminal regulatory landscape, destabilizing lineage identity without inducing complete lineage conversion. Together, our findings identify eGATA3 as a mechanistically distinct class of GATA3 mutation that promotes lineage plasticity through redistribution of genomic occupancy rather than loss of DNA-binding function alone.

## Introduction

Cell fate specification and maintenance are orchestrated by interconnected transcription factor networks that establish and sustain lineage-specific enhancer programs, thereby preserving cellular identity^1,2^. In breast cancer, disruption of these regulatory networks contributes to the emergence of distinct cellular states that can underlie clinically relevant molecular subtypes^3–6^. Breast cancer has traditionally been classified based on the expression of estrogen receptor (ER), progesterone receptor (PR), and human epidermal growth factor receptor 2 (HER2). Gene expression profiling has further refined these into intrinsic molecular subtypes, including luminal A, luminal B, HER2-enriched, basal-like, and normal-like^7–9^. Among these, luminal and basal-like tumors represent two major epithelial lineages and are distinguished by the activity of the GATA3–FOXA1–ESR1 transcription factor network, which is active in luminal tumors but largely inactive in basal-like tumors^10–14^. Consistent with this, the expression of GATA3, FOXA1, and ESR1 is highly correlated in luminal breast cancers^11,15^. Although these subtypes are histologically and clinically distinct, accumulating evidence suggests that luminal and basal-like states exist along a continuous transcriptional spectrum, highlighting an intrinsic capacity for lineage plasticity^16–18^. This plasticity has important clinical implications, as the transition toward a basal-like identity is frequently associated with more aggressive disease, therapeutic resistance, and poorer clinical outcomes^19,20^.

Increasing evidence suggests that perturbation of the GATA3–FOXA1–ESR1 transcription factor network contributes to lineage plasticity and endocrine therapy resistance in luminal breast cancer. Altered expression and recurrent missense mutations in ESR1 (Y537/D538) have been associated with the acquisition of basal-like features in luminal tumors^21,22^. Reduced expression of FOXA1 induces basal-like features^23^, but specific wing-domain FOXA1 mutations reinforce luminal identity, which confers endocrine therapy resistance^24,25^. Consistent with this concept, previous studies have also proposed that different classes of GATA3 mutations may exert distinct functional consequences depending on the affected protein domain^26^. Collectively, these observations demonstrate that disruption of lineage-defining transcription factor networks can produce diverse phenotypic outcomes depending on the molecular context and the nature of the underlying alteration.

Among the components of the luminal transcription factor network, GATA3 is a master regulator of luminal epithelial differentiation, and approximately 10% of the TCGA breast cancer cohort has mutations in GATA3^27–30^. In addition to genetic alterations, recent studies have shown that epigenetic reprogramming at GATA3-bound regulatory elements promotes lineage plasticity during the acquisition of endocrine therapy resistance, underscoring the central role of GATA3-mediated regulatory networks in maintaining luminal identity^31^. Protein-altering mutations in GATA3 include both frameshift and splice-site alterations with distinct structural consequences. Many truncating frameshifts and a recurrent splice-site mutation (neoGATA3) disrupt the second zinc-finger domain, impair DNA-binding activity, and rewire GATA3-mediated transcriptional control^11,32–35^. In contrast, extension mutations (eGATA3) arise downstream of both zinc-finger domains, generating proteins with an aberrant C-terminal peptide while retaining intact DNA-binding domains^36^ (**Fig. S1A**). This structural distinction, therefore, raises a fundamentally different mechanistic question of how a DNA-binding-competent mutant influences GATA3 function. Although previous studies have implicated eGATA3 variants in modulating therapeutic responses, including enhanced sensitivity to G9A/GLP inhibition^36^, their genome-wide chromatin-binding behavior, transcriptional regulatory capacity, and impact on luminal gene regulatory networks in hormone receptor-positive contexts remain largely unexplored.

Here, we investigate the functional consequences of eGATA3 mutations in luminal breast cancer. Combining the genomic, transcriptomic, and chromatin accessibility profiles of eGATA3-mutant patient tumors, we examine how the mutation reshapes GATA3 function. We further validated this by in vitro analysis of mutant GATA3 (P409) overexpressed in T47D cells and reanalysis of CRISPR knock-in MCF7 cells^37^. We find that eGATA3 retains chromatin association but alters the luminal regulatory landscape, leading to attenuation of luminal and estrogen-responsive transcriptional programs and progressive loss of luminal identity. Together, our findings identify eGATA3 as a mechanistically distinct class of GATA3 mutation that remodels the luminal regulatory landscape without abolishing DNA-binding capacity, thereby promoting lineage plasticity.

## Results

### eGATA3 mutations are associated with adverse clinical outcomes despite luminal characteristics

Extension frameshift mutations represent the predominant class of GATA3 alterations across primary (TCGA, SCANB) and metastatic (MSK-IMPACT) breast cancer cohorts (>55%; **Fig. S1B, C**), as compared to other mutation categories such as truncation and splice-site that result in loss of the second zinc-finger DNA-binding domain (**Fig. S1A)**. In the SCANB and MSK-IMPACT cohorts, the most frequent mutation was an extension frameshift at codons 408/409 (>40%; **Fig. 1A**), supporting the recurrent nature of these C-terminal GATA3 alterations (**Fig. S1D**). Interestingly, although the 408/409fs mutation represented the predominant hotspot, several independent frameshift mutations occurring at distinct positions generated extension proteins of remarkably similar length (**Fig. 1B,C**). This convergence suggests that selective pressure may act on the length of the C-terminal extension rather than on a specific mutation.

**Figure 1:**
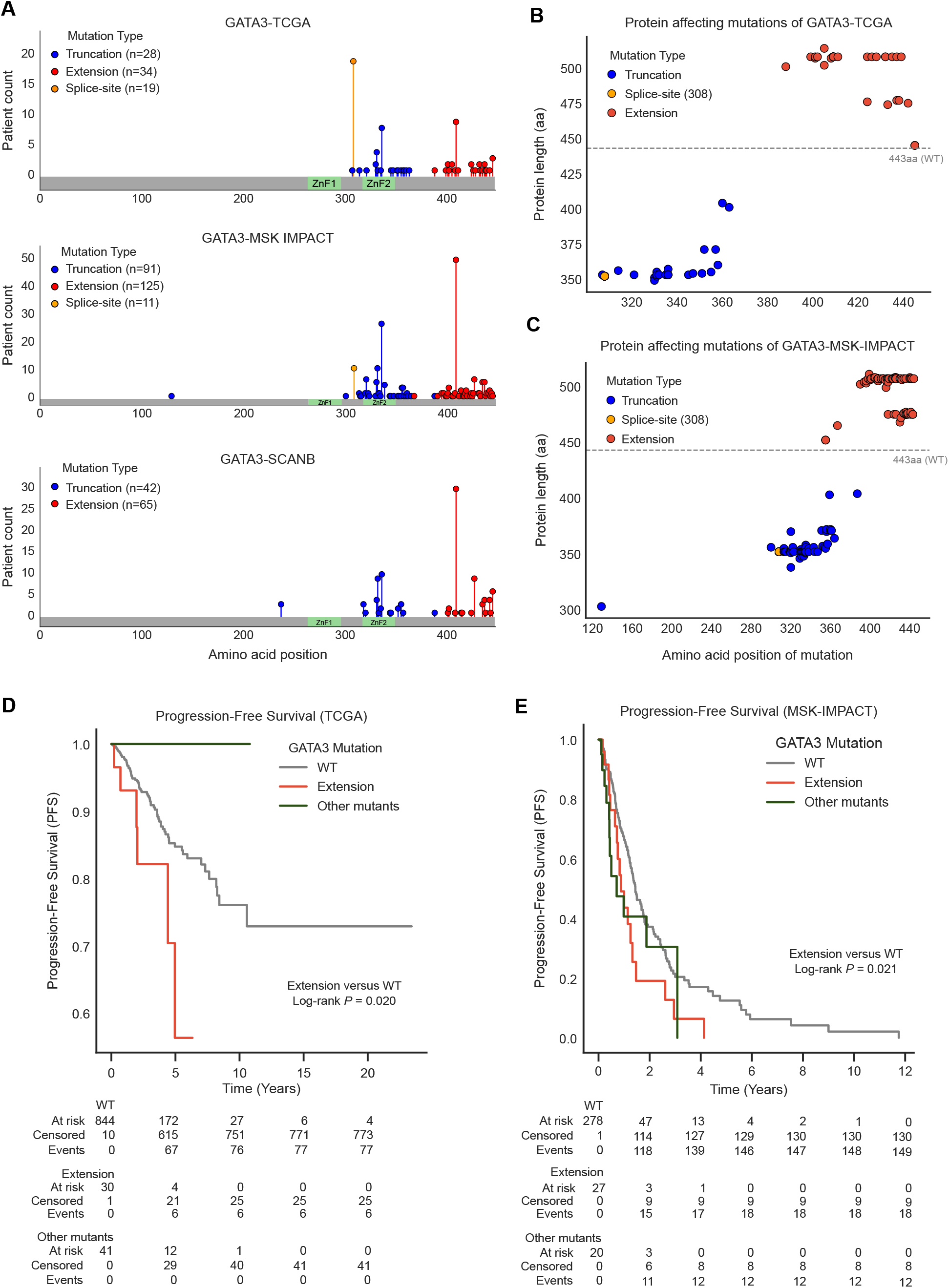
Clinical association of GATA3 mutations across primary and metastatic breast cancer cohorts. **(A)** Distribution of GATA3 mutation classes across the TCGA-BRCA, MSK-IMPACT, and SCAN-B breast cancer cohorts. **(B-C)** Consequence of frameshift and splice-site mutations on the protein sequence length in TCGA (B) and MSK-IMPACT (C). Each dot represents a GATA3 mutation in the patient and its corresponding predicted protein length. The dashed line indicates the length of wild-type GATA3 (443 amino acids). **(D–E)** Kaplan-Meier analyses of progression-free survival (PFS) in the TCGA-BRCA (D) and MSK-IMPACT (E) cohorts stratified by GATA3 mutation status. Statistical significance was assessed using a pairwise log-rank test comparing eGATA3-mutant and wild-type tumors. Other mutants comprise truncating and splice-site GATA3 mutants and are shown for comparison only.

We next examined the clinical association of eGATA3 mutations. Consistent with the established role of GATA3 in luminal lineage regulation, eGATA3 mutations were enriched in hormone receptor-positive luminal tumors (97% in TCGA and 89% in SCANB; **Fig. S2A**). Despite the luminal enrichment, patients with eGATA3 mutations had significantly shorter progression-free survival (PFS) than patients without GATA3 mutations (henceforth, wild-type) in both TCGA and MSK-IMPACT cohorts (Figs. 1D, E), with this association being more pronounced in the metastatic cohort (**Fig. S2B, C**). eGATA3-mutant patients also presented with multiple metastatic sites at a significantly higher frequency than wild-type controls (36.8% vs. 27.8%; **Fig. S2D**), indicative of a more disseminated metastatic phenotype in the MSK-IMPACT cohort. Analysis of other driver mutations with eGATA3 in the TCGA cohort showed no consistent co-occurrence with common breast cancer mutations (**Fig. S2E**), suggesting that the clinical features associated with eGATA3 are unlikely to be explained by concurrent canonical driver events.

Together, these analyses identify eGATA3 as a recurrent class of GATA3 mutations that are enriched in luminal tumors but are associated with shorter PFS and more extensive metastatic disease. Given these consistent clinical associations across independent patient cohorts, we next investigated the molecular consequences of eGATA3 in luminal breast cancer.

### eGATA3 reprograms the luminal transcriptome in breast cancer

To determine how distinct GATA3 mutations alter transcriptional programs, we analyzed RNA-seq data from TCGA breast cancers stratified by GATA3 mutation class (**Fig. S3A-C**). Differential expression analysis relative to GATA3 wild-type tumors identified distinct sets differentially expressed genes (FDR < 0.05, |log_2_FC| > 1) for truncating, splice-site, and eGATA3 mutants, with minimal overlap among the three mutation classes (**Fig. 2A**). Gene Ontology analysis further revealed enrichment of mutation class-specific biological processes (**Fig. 2B, C**). Among downregulated genes, eGATA3 mutants showed prominent enrichment for biological processes related to morphogenesis, cell adhesion, and extracellular matrix organization.

**Figure 2:**
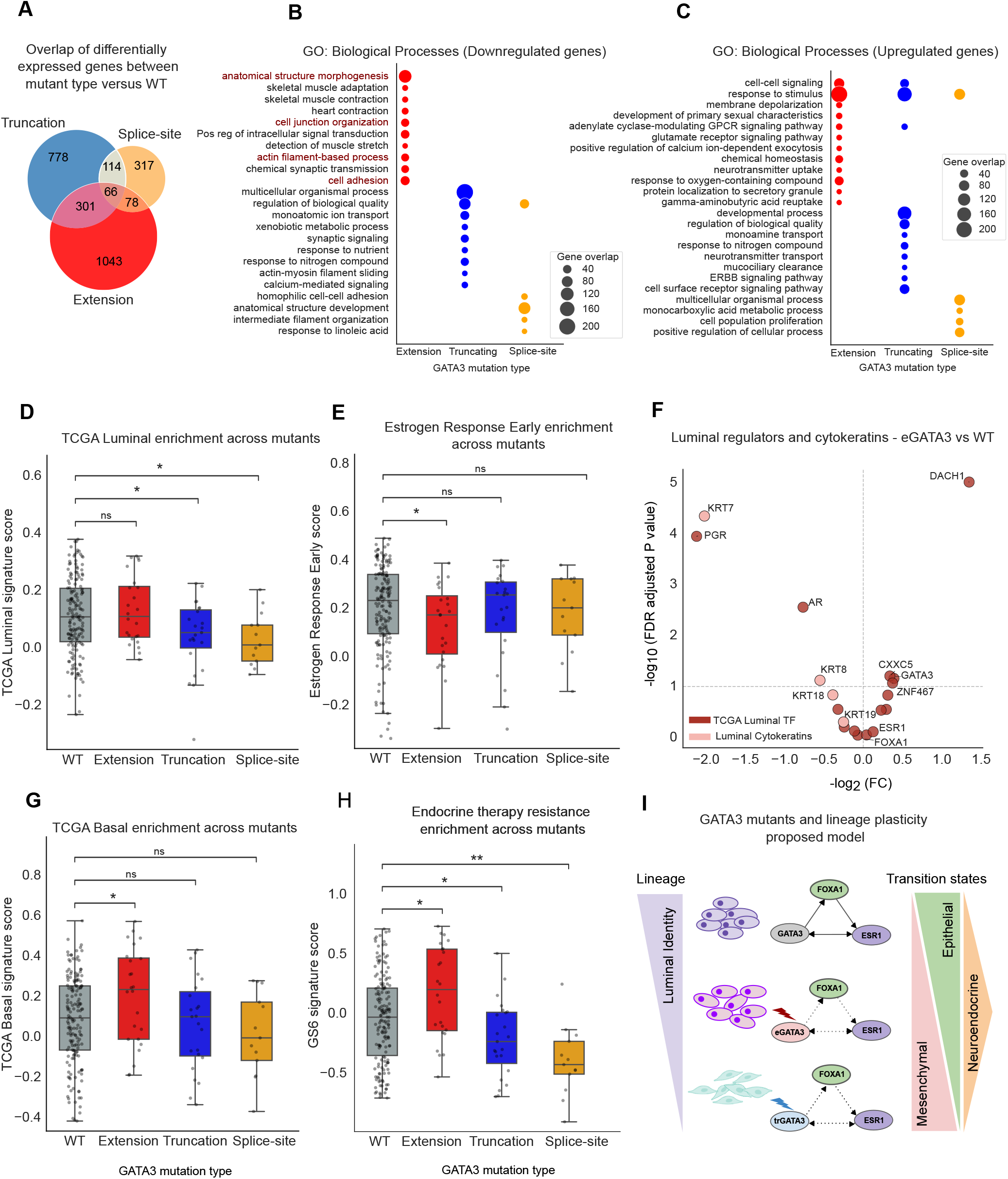
eGATA3 rewires the luminal transcriptome in breast tumors. **(A)** Venn diagram showing the overlap of differentially expressed genes (DEGs) among tumors harboring truncating, splice-site, and extension mutations relative to GATA3 wild-type tumors in the TCGA cohort. **(B-C)** Gene Ontology (GO) enrichment analysis of genes downregulated (B) and upregulated (C) in eGATA3-mutant tumors **(D-E);** GSVA scores for the TCGA luminal (D) and Estrogen Response Early (E) signatures across GATA3 mutants in TCGA breast tumors. **(F)** Differential expression analysis of cytokeratin genes and TCGA-derived lineage-associated TFs in eGATA3-mutant tumors with respect to WT tumors. Dashed lines indicate the differential expression thresholds used for significance. PGR, progesterone receptor; AR, androgen receptor; KRT7, keratin 7; KRT8, keratin 8; KRT18, keratin 18; KRT19, keratin 19. **(G-H)** GSVA scores for the TCGA basal (G) and endocrine therapy resistance (GS6) (H) signatures across GATA3 mutants in TCGA breast tumors. **(I)** Proposed model illustrating the effect of eGATA3 on luminal lineage identity. In panels D, E, and G, boxes denote the interquartile range (Q1–Q3), the central line indicates the median, and whiskers extend to 1.5× the interquartile range. Data points beyond the whiskers are displayed as outliers. Statistical significance was assessed using the Mann-Whitney U test (two-sided). GSVA scores of FDR < 0.05 were considered significant. Asterisks denote statistical significance: *P* < 0.05 (*), *P* < 0.01 (**), and *P* < 0.001 (***); ns, not significant.

As developmental and cell-adhesion pathways are closely linked to luminal differentiation, we next asked whether eGATA3 alters the luminal transcriptional program. To assess luminal lineage identity, we evaluated four independent luminal gene signatures derived from Sørlie et al., Huper et al., and the TCGA and METABRIC breast cancer cohorts (**Supplementary Table 1)**^16^. These signatures capture luminal identity across distinct cellular and patient-tumor contexts. Given that our analyses were performed in the TCGA-BRCA cohort, we used the TCGA-derived luminal signature for the primary analysis. Despite the dysregulation of developmental and cell-adhesion pathways, eGATA3-mutant tumors largely retained the global luminal differentiation (**Fig. 2D, Fig. S3D-F**). We hypothesized that, since eGATA3 retains both DNA-binding domains, the mutant might selectively remodel specific components of the luminal program, rather than disrupting the luminal program globally. We therefore examined hormone receptor signaling together with lineage-associated transcription factors, using the TCGA-derived lineage framework described previously^22^, and expression of luminal and basal cytokeratin markers (**Supplementary Table 1)**. Hallmark estrogen response early was significantly depleted in eGATA3-mutant tumors as compared to WT, whereas estrogen response late exhibited a similar trend (**Fig. 2E; Fig. S3G**). Consistent with reduced estrogen signaling, progesterone receptor signaling was also decreased in eGATA3 (**Fig. S3H**). We next examined the expression of lineage-defining transcription factors and cytokeratin markers. Despite attenuation of estrogen-responsive transcriptional programs, expression of the master luminal transcription factors ESR1 and FOXA1 remained largely unchanged in eGATA3 (**Fig. 2F).** In contrast, eGATA3-mutant tumors selectively downregulated progesterone receptor (PGR), a canonical ER target gene, androgen receptor (AR), a luminal lineage regulator, and the luminal cytokeratins KRT7, KRT8, KRT18, and KRT19 (**Fig. 2F**) as compared to the WT. Together, these analyses show that eGATA3 selectively rewires the luminal transcriptome.

We next asked whether the alteration in the luminal transcriptome was accompanied by the acquisition of basal-associated transcriptional features. Among the GATA3 mutation classes, only eGATA3 tumors exhibited significant enrichment of the TCGA basal gene signature as compared to WT (**Fig. 2G, Fig. S4A-D**). However, this enrichment was not accompanied by induction of canonical basal lineage transcription factors or basal cytokeratins (**Fig. S4E**), indicating that eGATA3 promotes basal-associated transcriptional programs without complete basal lineage conversion. Consistent with this observation, projection of GATA3-mutant tumors along with WT tumors (including luminal and basal-like) onto a UMAP generated using the TCGA basal gene signature showed that eGATA3 tumors remained within the luminal transcriptional landscape (**Fig. S4F**). Because disruption of luminal identity and lineage plasticity have been associated with endocrine therapy resistance^22,30,38^, we next examined whether eGATA3-mutant tumors exhibited transcriptional features of endocrine therapy resistance. The GS6 tamoxifen resistance signature was selectively enriched in eGATA3-mutant tumors (**Supplementary Table 1, Fig. 2H**), suggesting that eGATA3-mediated remodeling is accompanied by transcriptional programs associated with endocrine therapy resistance.

Acquisition of basal-associated transcriptional features is frequently associated with epithelial-mesenchymal transition (EMT), a transcriptional program linked to increased cellular plasticity^18,39,40^. We therefore asked whether eGATA3 tumors activated EMT-associated programs^39^. However, eGATA3 tumors showed no enrichment of epithelial, mesenchymal, or partial EMT gene signatures (**Supplementary Table 1, Fig. S5A).** Similarly, the expression of canonical EMT transcription factors remained largely unchanged, except that of TWIST1, which was upregulated (**Fig. S5B**). These findings indicate that the transcriptional reprogramming associated with eGATA3 occurs independent of a canonical EMT program.

In the absence of EMT activation, we investigated whether eGATA3 was associated with an alternative transcriptional program that is linked to lineage plasticity. Neuroendocrine differentiation has been described in a subset of epithelial cancers and has previously been associated with GATA3-mutant tumors^41–44^. We found that eGATA3 tumors exhibited significant upregulation of classical neuroendocrine marker genes (**Fig. S5C-E**). Furthermore, GSVA analysis using neuroendocrine prostate cancer-derived gene signatures^45^ demonstrated significant enrichment of neuroendocrine-associated transcriptional programs specifically in eGATA3 tumors (**Supplementary Table 1, Fig. S5F**). Together, these findings indicate that eGATA3-associated lineage plasticity is accompanied by activation of neuroendocrine-associated transcriptional features rather than a canonical EMT program.

Together, these findings indicate that eGATA3 does not cause a complete loss of luminal identity but instead establishes an altered transcriptional state characterized by attenuation of hormone-responsive and luminal programs together with acquisition of basal- and neuroendocrine-associated features (**Fig. 2I**). To determine how this altered transcriptional state arises, we next asked whether eGATA3 remodels the underlying chromatin landscape in patient tumors.

### eGATA3 is associated with the chromatin accessibility remodeling in breast tumors

To investigate whether eGATA3 remodels the chromatin landscape in patient tumors, we performed differential chromatin accessibility analysis using TCGA ATAC-seq data, comparing eGATA3-mutant (n = 4) and GATA3 wild-type tumors (n = 39). This identified 1,848 regions with increased accessibility and 1,401 regions with decreased accessibility in eGATA3 tumors (**Fig. 3A**). Genomic annotation showed that most differentially accessible regions were in distal intronic and intergenic regulatory elements, with relatively few changes at promoters (**Fig. S6A, B**), suggesting that eGATA3 might mainly remodel enhancer-associated chromatin.

**Figure 3:**
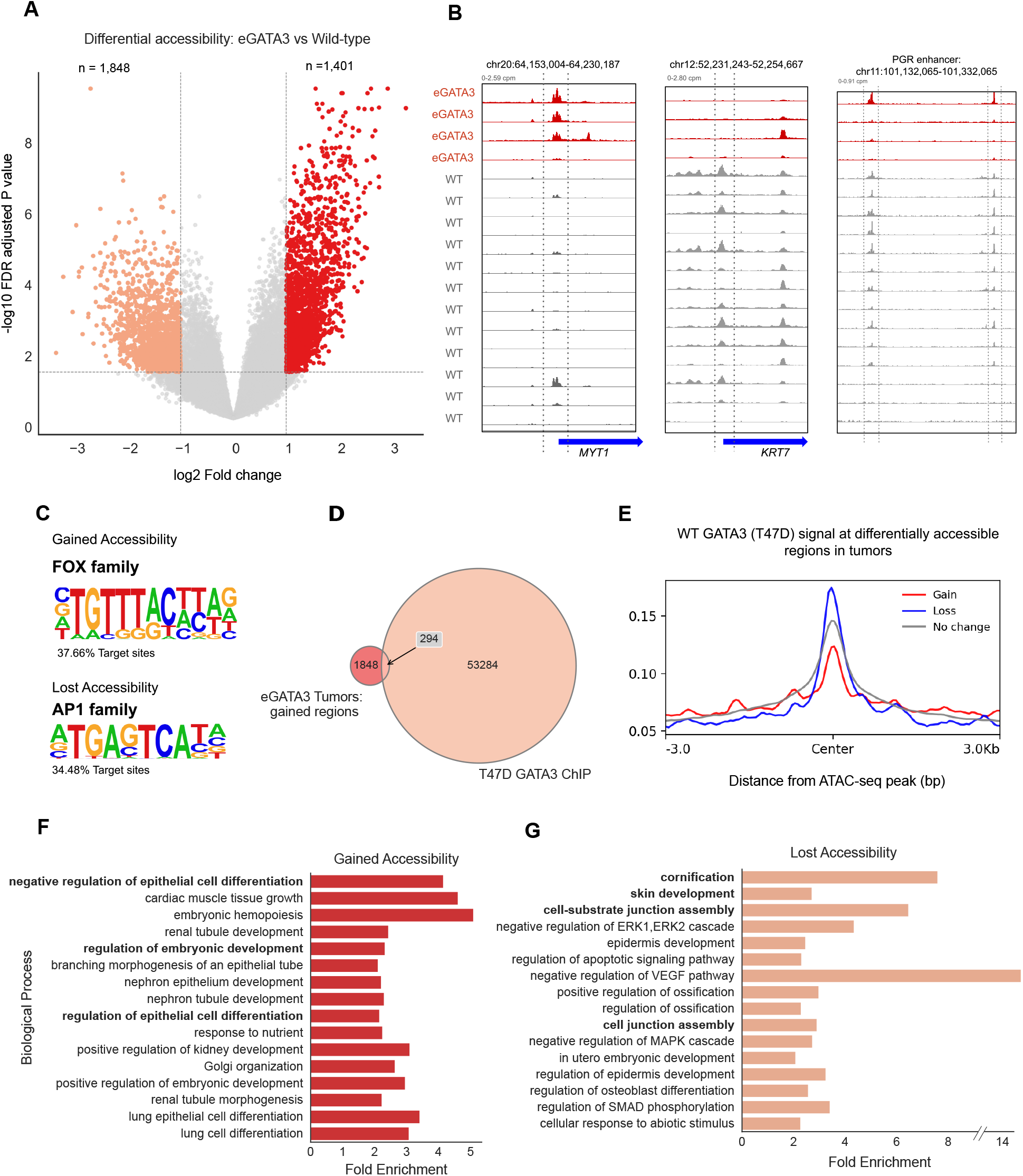
eGATA3 is associated with chromatin accessibility remodelling in breast tumors. **(A)** Differential chromatin accessibility analysis of eGATA3-mutant tumors relative to GATA3 wild-type tumors using TCGA ATAC-seq data. Differentially accessible regions are highlighted in red (gained accessibility) and orange (lost accessibility). **(B)** Representative genome browser tracks showing chromatin accessibility at the **MYT1** locus, **KRT7** locus, and **PGR** locus in eGATA3-mutant tumors. **(C)** Motif enrichment analysis of regions exhibiting increased accessibility identified significant enrichment of the FOX-family DNA-binding motif. Motif enrichment analysis of regions exhibiting decreased accessibility identified enrichment of the AP-1 family transcription factor motif. **(D)** Overlap between regions exhibiting increased chromatin accessibility in eGATA3-mutant tumors and GATA3 binding sites identified by ChIP-seq in T47D cells. **(E)** Aggregate wt-GATA3 ChIP-seq signal centered on regions with increased, decreased, or unchanged chromatin accessibility in eGATA3-mutant tumors. **(F)** GREAT analysis of genes associated with regions exhibiting increased chromatin accessibility. **(G)** GREAT analysis of genes associated with regions exhibiting decreased chromatin accessibility.

Although promoter-associated changes were less frequent, representative loci illustrated a close correspondence between chromatin accessibility and gene expression (**Fig. 3B; Fig. S6C, D**). Increased accessibility at the MYT1 promoter was associated with increased MYT1 expression; MYT1 is a transcriptional regulator implicated in neuroendocrine differentiation^46^. In contrast, the luminal epithelial marker KRT7 showed reduced promoter accessibility accompanied by decreased expression (**Fig. 3B; Fig. S6C,D**). Notably, distal regulatory elements associated with the hormone receptor gene PGR also showed reduced accessibility in eGATA3 tumors (**Fig. 3B**), consistent with reduced PGR expression and attenuated estrogen-responsive transcription (**Fig. 2E,F**). This regulatory region overlaps the PGR locus previously examined in ZnF2-truncating GATA3 mutants^34^, in which GATA3 binding was altered at multiple regulatory sites associated with PGR. Together, these findings indicate that chromatin remodeling in eGATA3 tumors accompanies selective erosion of luminal gene regulation alongside acquisition of alternative transcriptional features.

Because most accessibility changes occurred at distal regulatory elements, we next applied de novo motif analysis to identify transcription factors associated with these regions. Regions that gained accessibility were significantly enriched for FOX-family motifs, whereas those that lost accessibility were preferentially enriched for AP-1 family motifs (**Fig. 3C**). AP-1 family transcription factors can cooperate with ERα and GATA3 at regulatory elements, suggesting that the loss of accessibility at AP-1-associated regions may contribute to the attenuation of estrogen signaling observed above^11,47^. To further identify motifs that distinguish redistributed sites from baseline, we repeated motif enrichment analysis using non-changing accessibility peaks as the background, which similarly identified FOX-family motifs at gained sites and AP-1 motifs at lost sites (**Fig. S7A, B).**

Since GATA3 and FOXA1 are known to colocalize^11,48^, we next asked whether the FOX motif-enriched gained regions were associated with GATA3 binding. We compared the differentially accessible regions with published GATA3 ChIP-seq data from the T47D cells (which does not harbor any mutations in GATA3)^34^. Gained regions displayed relatively minimal overlap with canonical WT GATA3 binding sites and low WT GATA3 occupancy, whereas lost regions displayed substantially stronger WT GATA3 binding (Figs. 3D, E). Thus, gained regions in eGATA3 tumors are largely distinct from canonical GATA3-bound regulatory elements, despite the enrichment for FOX-family motifs. Together, these findings suggest that eGATA3-associated chromatin remodeling involves a shift away from regulatory elements normally occupied by WT GATA3 toward distinct, FOX-associated regulatory elements.

The altered chromatin accessibility observed in eGATA3 tumors could reflect impaired DNA recognition by the mutant protein or altered regulatory engagement despite preservation of DNA binding. To determine whether eGATA3 can still bind DNA, we predicted the structure of the eGATA3 dimer in complex with the GATA DNA motif using AlphaFold3 (AF3) (see Methods)^49^. The predicted structure for the wild-type GATA3 with DNA is similar to that of eGATA3 with DNA (TM-score of 0.9 from USalign)^50^. This indicates that the zinc-finger domain remains intact and capable of engaging DNA in a manner comparable to wild-type GATA3 (**Fig. S7C**). These findings suggest that the chromatin remodeling associated with eGATA3 is unlikely to result from impaired DNA-binding capacity. Rather, they support a model in which the intact DNA-binding domain enables an altered engagement of eGATA3 with the regulatory regions. This pattern is consistent with the possibility that the C-terminal extension affects cofactor interactions or transactivation capacity while preserving the structural integrity of the DNA-binding domain.

GREAT analysis for gene ontology of regions with gained accessibility showed enrichment of pathways linked to negative regulation of epithelial differentiation and embryonic development, while regions losing accessibility were enriched for epithelial structural programs, including cornification, cell junction organization, and skin development (**Fig. 3F, G**). Interestingly, regions losing accessibility were also enriched for pathways related to negative regulation of vascular endothelial growth factor (VEGF) signaling, raising the possibility that eGATA3-associated chromatin remodeling may contribute to the more disseminated metastatic phenotype observed in eGATA3-mutant tumors (**Fig. 3G**). Together, these enrichments suggest that eGATA3-associated chromatin remodeling weakens luminal epithelial identity and promotes a more developmentally plastic regulatory landscape. To investigate whether the chromatin and transcriptional changes observed in patient tumors are caused by eGATA3, we established a luminal breast cancer model to investigate the transcriptional consequences of the mutant in a controlled cellular context.

### eGATA3 over-expression induces luminal transcriptional reprogramming in breast cancer cell lines

To determine the eGATA3-driven transcriptional changes, we modeled the recurrent extension GATA3 mutation P409fs in luminal T47D breast cancer cells. Because GATA3 mutations are predominantly heterozygous in patients^34^, eGATA3 was stably expressed alongside endogenous wild-type GATA3 using two independent delivery methods, transfection and transduction. The P409fs mutation results in an extended protein by continuing translation into the downstream untranslated sequence present in the wild-type transcript; hence, the mutant and wild-type transcripts cannot be readily distinguished by standard RNA-expression assays. Therefore, mutant expression was confirmed at the protein level by Western blot (**Fig. 4A**). RNA sequencing confirmed that eGATA3 expression led to concordant depletion of luminal and estrogen-response programs in both transfected and transduced lines (**Fig. 4B, S8A**), demonstrating that the transcriptional changes seen in eGATA3 tumors are reproducible across experimental systems and are likely driven by eGATA3 itself. For simplicity, all subsequent analyses were conducted on the stably transfected line.

**Figure 4:**
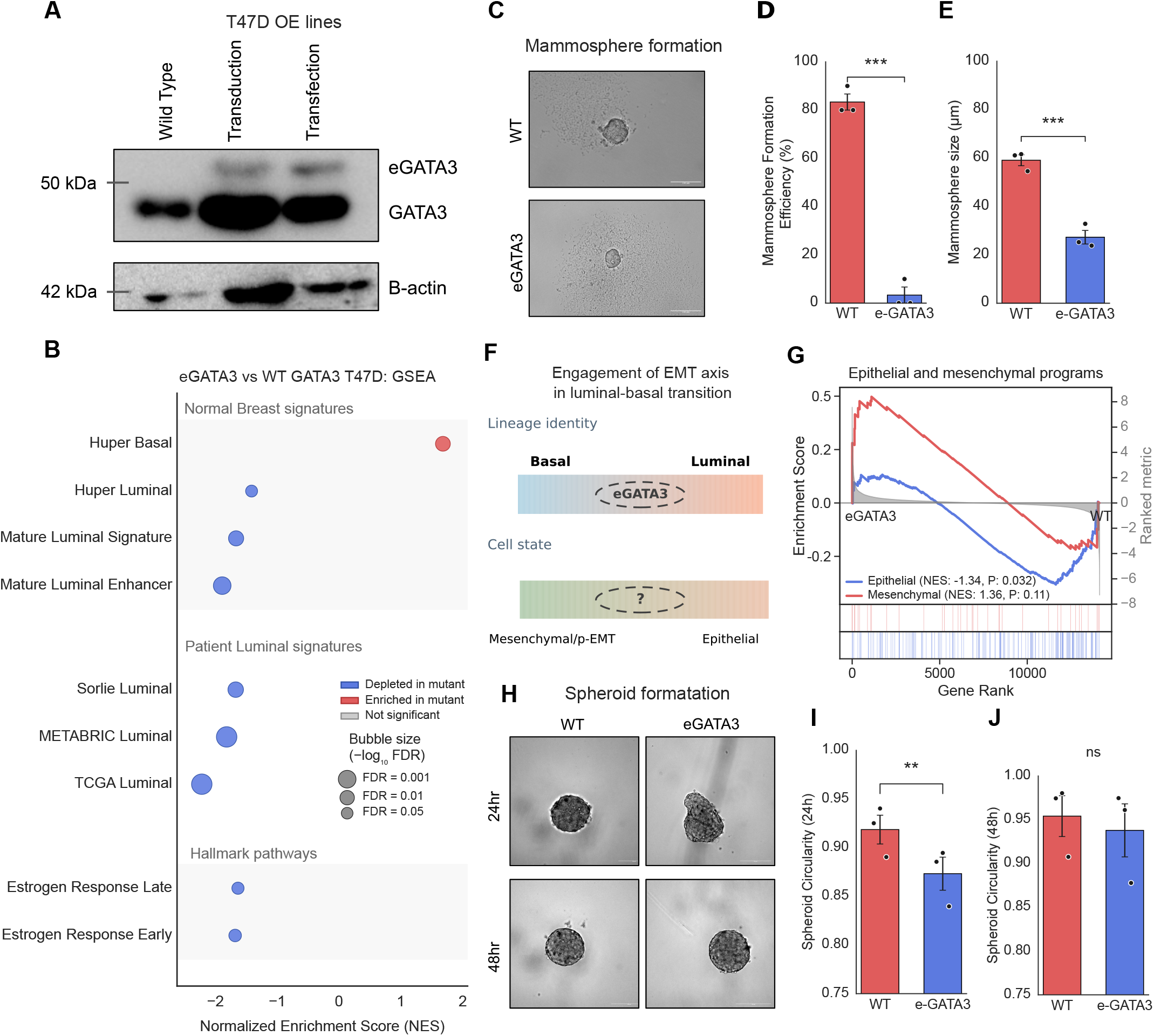
eGATA3 induces luminal transcriptional reprogramming and impairs epithelial organization in breast cancer cells. **(A)** Western blot analysis confirming expression of the eGATA3 P409fs mutant alongside endogenous wild-type GATA3 in stable T47D transfection and transduction models. **(B)** Gene set enrichment analysis (GSEA) of RNA-sequencing data from eGATA3-transfected T47D cells relative to wild-type cells. **(C)** Representative brightfield images of mammospheres formed by wild-type and eGATA3-expressing T47D cells. Scale bar 100 µm. **(D)** Quantification of mammosphere formation efficiency demonstrating reduced self-renewal capacity in eGATA3-expressing cells. **(E)** Quantification of mammosphere size showing reduced mammosphere diameter in eGATA3-expressing cells. **(F)** Schematic illustrating the coordinated loss of luminal lineage identity and epithelial cell state induced by eGATA3 expression. **(G)** GSEA analysis demonstrating depletion of an epithelial and mesenchymal gene signature in eGATA3-expressing cells. **(H)** Representative brightfield images of spheroids formed by wild-type and eGATA 3-expressing T47D cells at 24 and 48 hours. Scale bar 100 µm. **(I)** Quantification of spheroid circularity at 24 hours demonstrating impaired spheroid compaction in eGATA3-expressing cells. **(J)** Quantification of spheroid circularity at 48 hours showing no significant difference between wild-type and eGATA3-expressing cells. In panel D, E, I, J, data presented as mean ± S.E.M. Statistical significance was determined using a two-tailed Student’s *t-test.* Asterisks denote statistical significance: *P* < 0.05 (*), *P* < 0.01 (**), and *P* < 0.001 (***); ns, not significant.

In contrast to the more selective transcriptional rewiring observed in patient tumors, eGATA3 expression in T47D cells produced a broader attenuation of luminal lineage programs, with depletion of multiple independent luminal signatures, including normal mammary-derived (Huper luminal, mature luminal, and mature luminal enhancer) and tumor-derived (TCGA, METABRIC, and Sørlie) signatures (**Supplementary Table 1, Fig. 4B**). Reanalysis of a previously published ZnF2-truncating GATA3 mutant RNA-seq dataset^34^ showed similar attenuation of luminal transcriptional programs (**Fig. S8B**), indicating that both extension and ZnF2-truncating GATA3 mutants suppress luminal lineage programs despite their distinct structural consequences. Consistent with the depletion of the luminal program, the canonical luminal regulator PGR was downregulated (log₂FC = −0.57, FDR = 1.45 × 10⁻³; **Fig. S8C**), and the Hallmark Estrogen Response Early and Late signatures were also downregulated (**Fig. 4B)**. Gene Ontology analysis of differentially expressed genes further identified changes in developmental, cell adhesion, cell motility, and epithelial proliferation-associated processes in eGATA3-expressing cells (**Fig. S8C,D**). We also observed that gene sets associated with tamoxifen resistance in T47D^51^ were concurrently enriched (**Supplementary Table 1, Fig. S8E**), suggesting that eGATA3-mediated luminal reprogramming may contribute to resistance to endocrine therapy.

To assess whether functional changes accompanied the attenuation of luminal transcriptional programs, we performed mammosphere formation assays (see Methods) to determine the self-renewal and clonogenic potential of luminal progenitor-like cells in anchorage-independent conditions (**Fig. 4C**). The mammosphere formation efficiency of eGATA3-expressing cells was significantly decreased compared with WT cells (two-sided t-test, P < 0.001), indicating impairment of the sphere-initiating capacity. In addition, the mammospheres were much smaller (**Figs. 4D, E;** P < 0.001), indicating a decreased proliferative expansion of sphere-forming cells. Together, these results show that eGATA3 impairs functional properties related to luminal progenitor-like cells, in line with the general attenuation of luminal transcriptional programs seen in T47D cells.

Loss of luminal lineage identity has been closely linked to changes in epithelial cell state (**Fig. 4F**). Consistent with this relationship, depletion of luminal gene programs in eGATA3-expressing cells was accompanied by significant negative enrichment of an epithelial gene signature (**Fig. 4G**). Gene ontology analysis further revealed remodeling of cell adhesion pathways, including downregulation of epithelial adhesion genes such as CLDN1, CLDN16, and DAB2, and increased expression of mesenchymal adhesion-associated genes, including CD44 and MMP14 (**Fig. S8C**). Functional assessment using spheroid formation assays (see Methods) demonstrated delayed spheroid compaction and reduced circularity at 24 hrs, consistent with impaired cell-cell adhesion and disruption of epithelial organization (**Fig. 4H-J**). Despite these changes, eGATA3-expressing cells did not show increased migratory capacity in wound-healing assays (**Fig. S9A, B**), indicating that the disruption of epithelial organization does not lead to a fully mesenchymal phenotype. Nevertheless, collective dissemination does not require a mesenchymal migratory state, as studies have shown novel remodeling of adhesion can result in circulating tumor clusters with metastatic potential^52^. This possibility is particularly relevant given our observation that eGATA3-mutant tumors were more frequently associated with multiple metastatic sites at diagnosis, raising the possibility that altered epithelial organization may contribute to this.

To determine whether eGATA3-induced phenotypic changes were accompanied by altered proliferation, we performed colony formation assays. eGATA3 expression did not significantly affect colony number, although colonies were modestly smaller than those formed by wild-type cells, consistent with impaired cell-cell adhesion (**Fig. S9C**). These findings suggest that eGATA3-induced transcriptional reprogramming does not reflect changes in proliferation.

Finally, we asked how closely the T47D model recapitulates the transcriptional alterations observed in patient tumors at the gene level. Genes downregulated in eGATA3-mutant tumors were significantly enriched among genes downregulated in eGATA3-expressing T47D cells (NES = −2.171, P < 0.0001), whereas genes upregulated in patient tumors showed a similar but weaker positive trend (NES = 1.224, *P* = 0.07) (**Fig. S9D**). In contrast to patient tumors, eGATA3-expressing T47D cells did not exhibit enrichment of tumor-derived basal transcriptional signatures (**Fig. S9E**). This suggests that while destabilization of the luminal program represents a cell-autonomous consequence of eGATA3 expression, acquisition of the basal-associated transcriptional features observed in patient tumors likely requires additional tumor-specific or microenvironmental cues.

Collectively, these findings demonstrate that eGATA3 directly destabilizes the luminal transcriptional program and associated epithelial organization, without inducing corresponding basal-like or mesenchymal characteristics, suggesting a state of weakened luminal identity rather than complete lineage conversion. We next investigated how eGATA3 reshapes GATA3 chromatin occupancy to drive this transcriptional reprogramming.

### eGATA3 alters binding occupancy in breast cancer cell lines

The differential chromatin accessibility observed in eGATA3-mutant tumors, together with the preferential loss and gain of accessibility at distinct classes of regulatory elements including FOX and AP1 families, suggested altered engagement of regulatory regions in the mutants. To determine whether this altered chromatin engagement is a direct consequence of eGATA3, we next profiled GATA3 chromatin occupancy using CUT&RUN in stably eGATA3-expressing T47D cells described above (see Methods). Comparison of GATA3 binding revealed extensive redistribution, with 7,988 sites unique to parental cells, 18,862 unique to eGATA3-expressing cells and 14,766 shared between the two conditions (**Fig. 5A**). Analysis of GATA3 occupancy across these three peak classes confirmed strong binding at shared sites and distinct occupancy patterns at WT- and eGATA3-unique regions (**Fig. 5B**).

**Figure 5:**
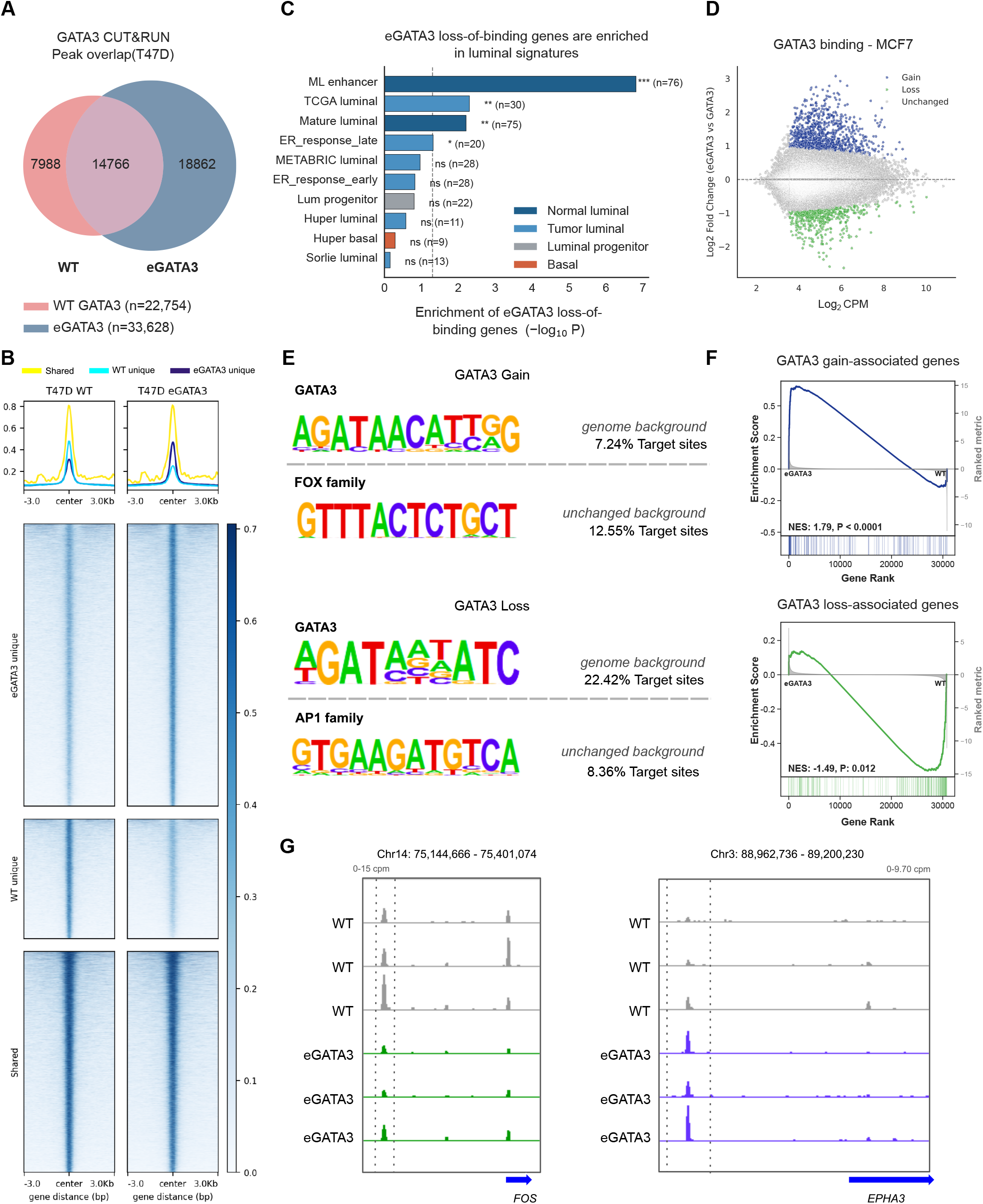
eGATA3 alters binding patterns and luminal regulatory landscape. **(A)** Venn diagram showing shared and condition-specific GATA3 CUT&RUN peaks in wild-type and eGATA3-expressing T47D cells. **(B)** Average GATA3 CUT&RUN signal at wild-type-unique and eGATA3-unique binding sites. **(C)** Luminal enrichment of genes associated with wild-type-unique GATA3 binding sites using normal mammary-derived and tumor-derived luminal gene signatures. **(D)** Differential GATA3 binding analysis using an independent GATA3-MCF7 ChIP-seq dataset. **(E)** De novo motif enrichment analysis of regions with gain and loss in GATA3 occupancy with genome background and unchanged regions as background. **(F)** Gene set enrichment analysis demonstrating that genes associated with gain in GATA3 binding are enriched among upregulated genes, whereas genes associated with loss in GATA3 binding are enriched among downregulated genes. **(G)** Genome browser tracks illustrating loss and gain of GATA3 occupancy at representative loci following eGATA3 expression.

To determine whether the observed redistribution contributes to the loss of luminal identity, we annotated genes associated with sites that lost GATA3 occupancy (WT-unique peaks). Overall, the genes that lost GATA3 binding were significantly downregulated in eGATA3-T47D cells (promoter-associated genes, NES = −1.35, P = 0.02; enhancer-associated genes, NES = −1.59, P < 0.0001; **Fig. S10A,B**). Further, we assessed the genes associated with WT unique sites specifically for enrichment across the luminal gene signatures and found that they were represented in both normal mammary-derived and tumor-derived luminal signatures (**Fig. 5C**). Reduced GATA3 occupancy was observed at representative luminal regulators such as NRF2 (**Fig. S10C**). These findings indicate that loss of GATA3 binding occurs at luminal regulatory elements and provides a mechanistic basis for the transcriptional erosion of luminal identity.

Having seen that eGATA3 is associated with loss of GATA3 occupancy at luminal regulatory elements, we next examined whether there is also gain in occupancy at distinct regulatory regions. Ectopic expression increases total GATA3 abundance, so we cannot clearly attribute the increased occupancy seen in the T47D overexpression model to mutation-specific redistribution. To separate redistribution from increased GATA3 dosage, we analyzed published GATA3 ChIP-seq data from a CRISPR knock-in MCF7 model expressing the same P409fs allele from the endogenous locus^37^. Differential binding analysis identified 980 gained and 562 lost sites relative to unchanged regions (**Fig. 5D; Fig. S10D**).

To identify the sequence features associated with redistributed GATA3 binding in the MCF7 eGATA3 model, we performed de novo motif enrichment analysis of the ChIP-seq peaks. Relative to genomic background, GATA motifs were enriched at both gained and lost binding sites, confirming that both classes represent bona fide GATA3 regulatory elements. However, GATA motifs were more strongly enriched at lost sites (22% of targets) than at gained sites (7% of targets), suggesting that GATA3 binding is preferentially lost from canonical GATA-rich regulatory elements (**Fig. 5E**). To identify features distinguishing redistributed sites from stable GATA3 binding, motif enrichment was repeated using unchanged GATA3 peaks as the background. Under these conditions, lost sites were preferentially enriched for AP-1 family motifs (8.4% of targets), whereas gain sites were enriched for FOX-family motifs, predominantly FOXA1 (12.6% of targets) (**Fig**. **5E**). Consistent with these motif enrichments, gained GATA3 binding sites exhibited increased FOXA1 and ER occupancy, whereas lost sites showed reduced FOXA1 and ER occupancy in MCF7 eGATA3 cells (**Fig**. **S10F**). Together, these findings indicate that GATA3 redistribution occurs selectively across distinct classes of luminal regulatory elements, with binding preferentially lost at AP-1-enriched sites and gained at FOXA1- and ER-associated regulatory elements. This selective redistribution is consistent with the altered estrogen-response transcriptional programs observed in eGATA3 tumors (**Fig**. **2E**).

Finally, we asked whether redistribution of GATA3 occupancy was associated with altered transcription. Genes proximal to gained GATA3 binding sites were significantly enriched among genes upregulated in the MCF7 knock-in transcriptome (NES = 1.79, P < 0.0001), whereas genes associated with lost binding sites were enriched among downregulated genes (NES = −1.45, P = 0.012; **Fig. 5F**). Representative loci, including the AP-1 family member, FOS, and the receptor tyrosine kinase EPHA3, illustrate these occupancy changes (**Fig. 5G**). EPHA3 has previously been implicated in breast cancer cell motility and invasion^53^. Similarly, genes associated with lost GATA3 binding in MCF7 were significantly depleted in the T47D transcriptome also (NES = −1.44, P = 0.02; **Fig. S10E**), indicating that loss of GATA3 occupancy consistently predicts transcriptional repression across independent models. Together, these findings show that selective redistribution of GATA3 occupancy across distinct classes of luminal regulatory elements is linked to coordinated changes in gene expression, connecting altered chromatin occupancy to the transcriptional rewiring induced by eGATA3.

### eGATA3 drives progressive transcriptional and chromatin reprogramming at single-cell resolution

Having observed that eGATA3 redistributes occupancy across distinct classes of regulatory elements, we next asked how these changes manifest at single-cell resolution. We therefore performed paired single-nucleus RNA sequencing (snRNA-seq) and ATAC-sequencing (snATAC-seq) of wild-type and eGATA3-expressing T47D cells (see Methods). UMAP visualization of the transcriptomic data showed clear separation of wild-type and eGATA3 cells, indicating distinct mutant-associated transcriptional states (**Fig. 6A, Figs. S11A, B**). Consistent with the bulk RNA sequencing analyses, mature luminal, epithelial and estrogen-response signatures along with the previously used luminal signatures from Huper, Sorlie, TCGA, and METABRIC studies were significantly attenuated in eGATA3 cells (**Figs. 6B-D; S11C-F),** confirming widespread destabilization of luminal identity at single-cell resolution.

**Figure 6:**
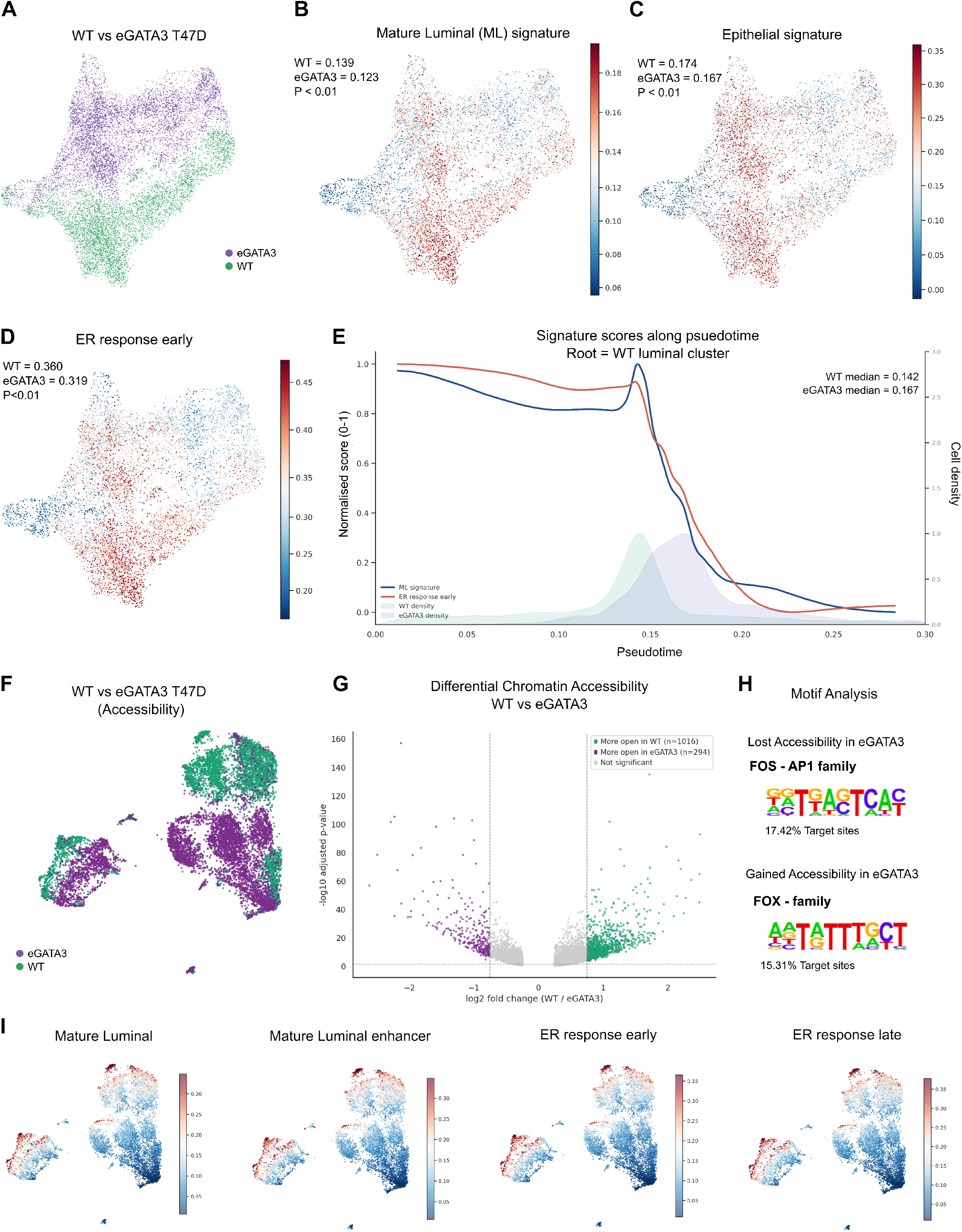
Single-nuclei multiomic profiling reveals transcriptional and chromatin reprogramming driven by eGATA3. **(A)** UMAP projection of single-nucleus RNA sequencing (snRNA-seq) profiles from wild-type and eGATA3-expressing T47D cells. **(B)** Projection of a mature luminal gene signature onto the snRNA-seq UMAP, showing reduced luminal program activity in eGATA3-expressing cells. **(C)** Projection of an estrogen-response early gene signature onto the snRNA-seq UMAP, demonstrating attenuation of estrogen-responsive transcription in eGATA3-expressing cells. **(D)** Projection of an epithelial gene signature onto the snRNA-seq UMAP, showing reduced epithelial program activity in eGATA3-expressing cells. **(E)** Diffusion Pseudotime analysis of wild-type and eGATA3-expressing cells, rooted in a WT luminal cell, showing progressive decline of mature luminal and estrogen-response signatures along the inferred trajectory. **(F)** UMAP projection of single-nucleus ATAC-sequencing (snATAC-seq) profiles from wild-type and eGATA3-expressing T47D cells. **(G)** Volcano plot showing differentially accessible chromatin regions between wild-type and eGATA3-expressing cells. **(H)** De novo motif enrichment analysis of regions exhibiting decreased or increased chromatin accessibility in eGATA3-expressing cells, showing enrichment of AP-1 family motifs in regions losing accessibility and FOX-family motifs in regions gaining accessibility. **(I)** Chromatin accessibility projected onto the snATAC-seq UMAP for mature luminal gene signatures, mature luminal enhancers, estrogen-response gene signatures early and late, demonstrating coordinated reduction in accessibility of luminal regulatory programs in eGATA3-expressing cells. Statistical significance was assessed using the two-sided Mann–Whitney U test.

Lineage transitions are often part of a continuous transcriptional process, and to determine whether the changes above reflected a progressive cell state transition, we reconstructed trajectories using Diffusion Pseudotime (DPT) (see Methods). To minimize the influence of proliferative states on trajectory inference, cells assigned to the G2/M phase were excluded prior to DPT analysis. Cells were distributed along a continuum, with a WT luminal cell selected as the root. Along this trajectory, both mature luminal and estrogen-response signatures progressively declined, suggesting gradual erosion of luminal identity (**Fig. 6E**). Concomitant with the decline in luminal identity, basal-like and neuroendocrine-associated signatures increased along DPT, indicating the acquisition of alternative lineage programs (**Fig. S12C**). While enrichment of these alternative lineage programs was modest at the population level (**Fig. S12A, B**), DPT analysis revealed their progressive emergence along the mutant-associated trajectory. eGATA3 cells were significantly enriched at later DPT states, corroborating the existence of a mutant-associated transcriptional trajectory.

We next asked whether the altered GATA3 occupancy observed previously was accompanied by changes in chromatin accessibility. We analyzed paired snATAC-seq profiles from WT and eGATA3-expressing T47D cells. UMAP projection showed clear differences in the chromatin accessibility landscape between the two conditions (**Fig. 6F**). Differential accessibility analysis identified both gains and losses of accessibility across the genome (**Fig. 6G**). Motif enrichment analysis of differentially accessible regions revealed that regions losing accessibility were enriched for AP-1 family motifs, whereas regions gaining chromatin accessibility were enriched for FOX family motifs, thus validating our earlier findings (**Fig. 6H**). This further suggests that changes in chromatin accessibility occur preferentially at distinct classes of regulatory elements following eGATA3 expression.

To determine the functional consequences of these chromatin accessibility changes, we quantified accessibility across regulatory regions associated with mature luminal enhancers, luminal lineage programs, and estrogen-responsive transcription. Accessibility across each of these regulatory programs was significantly reduced in eGATA3 cells compared to wild-type cells (**Fig. 6I**), which mirrored the transcriptional depletion of luminal and estrogen-response signatures seen in both bulk and single-cell RNA sequencing. Collectively, these results link changes in chromatin accessibility to the coordinated downregulation of luminal regulatory programs in eGATA3-expressing T47D cells.

Together with the GATA3 occupancy analyses, these findings support a model in which selective redistribution of GATA3 binding is associated with reorganization of the luminal regulatory network and coordinated changes in luminal gene expression.

## Discussion

GATA3 is among the most frequently mutated transcription factors in luminal breast cancer, yet the biological consequences of individual mutation classes remain incompletely understood. Here, we identify C-terminal extension mutations (eGATA3) as a mechanistically distinct class of GATA3 alteration that retains widespread chromatin occupancy while selectively redistributing GATA3 binding across the genome.

We found that the principal consequence of eGATA3 is attenuation of the luminal transcriptional program rather than direct specification of an alternative lineage state. Across patient tumors, eGATA3 reduced estrogen-responsive transcriptional programs, including expression of key luminal regulators such as PGR and AR. While other GATA3 mutants were associated with a more global loss of luminal identity, eGATA3 uniquely combined attenuation of luminal programs with enrichment of basal-associated transcriptional signatures in patient tumors. A similar coexistence of luminal and basal features has recently been described in the Y537S and D538G ESR1-mutant breast cancers, suggesting that disruption of the GATA3–FOXA1–ESR1 regulatory network can generate intermediate lineage states^16^. However, the underlying mechanisms appear fundamentally different. The ESR1 mutants maintain constitutive binding at ER target genes like PGR, whose hyperactivation directly leads to the expression of basal cytokeratins. While basalness is a direct result of the ESR1 mutation at PGR locus, our experimental models indicate that suppression of luminal identity is the primary consequence of eGATA3 expression. The basal-like features observed in patient tumors were not fully reproduced in vitro, raising the possibility that acquisition of alternative lineage programs requires additional microenvironment-dependent cues. Together, these observations suggest that distinct perturbations within the luminal lineage-specifying network can converge on similar transcriptional phenotypes through different molecular mechanisms.

Comparison with other GATA3 mutants further highlights that the disruption of luminal identity does not inevitably lead to the same alternative lineages and cell-states. Although eGATA3- and ZnF2-truncating mutants are structurally distinct; we observed that both exhibit attenuation of luminal transcriptional programs and reduced estrogen-responsive signalling. Consistent with previous reports, ZnF2 mutants also failed to confer a proliferative advantage in T47D cells in vitro, a phenotype similarly observed in our eGATA3 models^34^. Despite this transcriptional convergence, the downstream lineage trajectories differed. The previous study showed that ZnF2 mutants preferentially activate mesenchymal transcriptional programs, while eGATA3 primarily attenuates epithelial programs without inducing canonical EMT signatures or increasing migratory behavior. In addition, eGATA3-mutant patient tumors showed increased neuroendocrine-associated transcriptional programs, suggesting that loss of luminal identity may create a permissive state for multiple lineage trajectories depending on the mutational context. While the functional significance of these neuroendocrine features remains unknown, these observations suggest that different classes of GATA3 mutations converge in destabilizing the luminal regulatory network but lead to different cell states or lineage identities.

To investigate how eGATA3 attenuates luminal identity, we integrated chromatin accessibility profiling with CUT&RUN analysis of GATA3 occupancy. Our analysis showed that redistribution occurred across distinct classes of regulatory elements, with GATA3 binding preferentially lost at AP-1-enriched regions and gained at FOX-family-associated regulatory elements. Recent work has identified AP-1 as a collaborating transcription factor at productive GATA3 regulatory sites, where GATA3 and AP-1 frequently colocalize and contribute to productive enhancer formation^47^. This suggests that the preferential loss of GATA3 occupancy at AP-1-associated regions in eGATA3-expressing cells may reflect altered engagement with regulatory elements that normally support productive GATA3 activity. On the other hand, gained GATA3 occupancy was preferentially associated with FOX-enriched regulatory elements, along with increased FOXA1 and ER occupancy, suggesting that eGATA3 may redistribute GATA3 to a distinct class of luminal regulatory elements. Together, these findings suggest that preservation of the GATA3 DNA-binding domain is insufficient to maintain its normal regulatory function. Conversely, eGATA3 remains substantially associated with chromatin but redistributes its occupancy across different regulatory contexts, possibly modifying the regulatory architecture that preserves luminal identity.

The lineage plasticity associated with eGATA3 has important implications for endocrine therapy. We observed the enrichment of tamoxifen resistance signatures in eGATA3 mutant patient tumors and cell lines. ESR1 mutants and endocrine-resistant breast cancers have been shown to give rise to resistance through lineage plasticity^16,22^. Moreover, redistribution of GATA3 occupancy resembles the enhancer reprogramming reported during acquisition of endocrine resistance^5^. Given the loss of luminalness, the attenuation of ER signaling, and the redistribution of GATA3 binding sites in eGATA3 cells, we need to explore how ER-targeted therapies affect eGATA3 tumors. Although these observations do not establish functional endocrine resistance, they raise the possibility that eGATA3 creates a permissive regulatory state that facilitates adaptation to ER-targeted therapies. Apart from endocrine therapies, recent studies have reported selective sensitivity of eGATA3-expressing cells to G9a/GLP inhibition in non-malignant systems^36^. Whether this therapeutic vulnerability extends to hormone receptor-positive breast cancer remains an important question for future investigation.

Though our findings suggest selective redistribution of GATA3 occupancy as a defining feature of eGATA3, the molecular basis for this redistribution remains unknown. The C-terminal region of GATA3 has been shown to mediate protein-protein interactions^54^. Accordingly, the extended C-terminus of eGATA3 might change interactions with lineage-defining cofactors and redirect GATA3 occupancy to different regulatory elements. Consistent with this idea, regions exhibiting altered occupancy were enriched for motifs associated with FOX and AP-1 family transcription factors, both of which are known to cooperate with GATA3 in regulating luminal gene expression^10,11,47^. An alternative possibility is that the C-terminal extension influences higher-order transcriptional organization. Recent studies have shown that ZnF2-truncating GATA3 mutants impair transcriptional condensate formation and attenuate ER signaling by altering phase-separation dynamics^55^. The convergence of luminal loss across the mutants raises the possibility of a similar alteration in phase-separation dynamics in eGATA3 mutants, which requires future investigation.

The following are some of the limitations of our study. Our T47D overexpression model increases eGATA3 in the presence of endogenous GATA3, making it difficult to distinguish mutation-specific effects from consequences of altered GATA3 dosage. We therefore complemented these experiments with analyses of a CRISPR knock-in MCF7 model expressing the same P409fs allele from the endogenous locus. Nevertheless, the number of eGATA3-mutant patient tumors available for chromatin-accessibility analysis remains limited, and larger cohorts will be required to establish the extent and clinical relevance of these regulatory changes in vivo. In addition, our cell-line models recapitulate key features of eGATA3-driven transcriptional and chromatin remodeling, but they cannot fully capture the complexity of the tumor microenvironment, which is likely to influence lineage plasticity in vivo. Future studies using patient-derived organoids and animal models will be important to define how eGATA3 drives luminal reprogramming.

## Methods

### Patient cohorts and mutation classification

Protein-affecting GATA3 mutations were obtained independently from three breast cancer cohorts: The Cancer Genome Atlas (TCGA) Pan-Cancer Atlas (MC3, mc3.v0.2.8.PUBLIC.maf.gz)^56^, the MSK-IMPACT breast cancer cohort accessed through cBioPortal^57–59^, and the SCAN-B breast cancer cohort^60^. Mutation frequencies and mutation-class distributions were analyzed separately for each cohort. For TCGA, level 3 RNA sequencing data (Illumina HiSeq RNASeqV2, RSEM-normalized expression), somatic mutation data (MC3), copy-number alteration data (GISTIC2)^61^, and corresponding clinical annotations were obtained from the TCGA Pan-Cancer Atlas. Samples flagged for quality-control issues were excluded according to the TCGA Pan-Cancer quality annotations, and for patients with multiple tumor samples, only the primary tumor sample was retained for downstream analyses. Frameshift mutations were classified as extension or truncating based on the Human Genome Variation Society protein (HGVSp) annotation in the MC3 MAF file, which shows the position of the frameshift and the predicted change to the encoded protein. For transcriptomic and chromatin accessibility analyses, samples harbouring GATA3 copy-number alterations or TP53 mutations were excluded to minimise potential confounding effects. Unless otherwise stated, all downstream transcriptomic and chromatin accessibility analyses were performed using the TCGA cohort.

### Survival analysis

Clinical annotations for TCGA breast cancer patients were obtained from the TCGA Pan-Cancer Clinical Data Resource (TCGA-CDR)^62^. Progression-free survival (PFS) was used as the primary clinical endpoint for breast cancer, as recommended by the TCGA Pan-Cancer Atlas. Kaplan-Meier survival curves were generated using the Python lifelines package (version 0.30.0)^63^, and statistical significance between the groups was assessed using the paired log-rank test.

### Differential gene expression analysis of TCGA tumors

Raw gene-level read counts for TCGA breast cancer samples were obtained using the TCGAbiolinks R package (2.34.1)^64^. Protein-coding genes with low expression were filtered using the TCGAanalyze_Filtering function (method = quantile, qnt.cut = 0.25), and count data were normalized using TCGAanalyze_Normalization. Differential gene expression analysis between GATA3 mutation classes and wild-type tumors was performed using the TCGAanalyze_DEA function with the edgeR generalized linear model likelihood ratio test (glmLRT).

### Gene set and pathway enrichment analyses

Pathway enrichment analyses were performed on both patient and cell line transcriptomic datasets. For TCGA breast cancer samples, single-sample gene set variation analysis (GSVA) was performed using the GSVA^65^ R (2.0.7) package. Published luminal, basal, epithelial, neuroendocrine, and endocrine resistance gene signatures, together with MSigDB^66^ Hallmark gene sets, were used throughout the study as indicated in the corresponding analyses (Supplementary Table 1 and 2). For bulk RNA sequencing of T47D cells, preranked gene set enrichment analysis (GSEA) was performed using GSEApy (version 1.1.13)^67^. Gene ontology enrichment analysis of differentially expressed genes was performed using g:Profiler^68^, and significantly enriched biological processes were identified using a Benjamini-Hochberg-adjusted *P value* < 0.05.

### Differential chromatin accessibility analysis

ATAC-seq data for primary breast tumors were obtained from the TCGA breast cancer ATAC-seq dataset generated by Corces and Granja *et al*^69^. Raw peak count matrices were downloaded, and samples harboring eGATA3 mutations and GATA3 wild-type tumors were selected for downstream analyses. Technical replicates were retained throughout the analysis. Peaks with low accessibility were filtered by retaining regions with a minimum count of 1 count per million (CPM) in at least one sample using the edgeR package^70^ (4.4.2). Raw counts were transformed to log2(CPM) values with a prior count of 5 and normalized using the voom method. Differential chromatin accessibility analysis was performed using the limma^71^ package with empirical Bayes moderation (eBayes). Regions with an adjusted *P value* < 0.05 were considered differentially accessible.

### Genomic annotation and motif enrichment analysis

Differentially accessible regions were annotated to genomic features using HOMER^72^ (annotatePeaks.pl) with the hg38 reference genome. *De-novo* motif enrichment analysis was performed using HOMER (findMotifsGenome.pl). Unless otherwise stated, regions that did not exhibit significant changes in accessibility or occupancy were used as the background set for motif enrichment analyses. Gene ontology analysis of genes associated with differentially accessible regions was performed using the Genomic Regions Enrichment of Annotations Tool (GREAT) with default parameters ^73^.

### Cell culture and generation of stable eGATA3 cell lines

The human luminal breast cancer cell line T47D was obtained from the American Type Culture Collection (ATCC) and maintained in RPMI 1640 Medium (ATCC modification, Gibco, Cat. No. A1049101) supplemented with 10% fetal bovine serum (FBS; Gibco, Invitrogen #16000044) and 1% penicillin–streptomycin (Gibco, Invitrogen #15140163) at 37°C in a humidified incubator with 5% CO₂.The wild-type GATA3 expression construct TFORF2240 (Addgene plasmid #141984; RRID:Addgene_141984), a gift from Feng Zhang, was used as the template to generate the GATA3 P409fs extension mutant^74^. The DNA sequence encoding the mutant C-terminal extension was synthesized by GeneArt (Thermo Fisher Scientific, **Supplementary Table 3**) and cloned in-frame into the wild-type GATA3 construct using BamH1-HF (NEB #R3136) and Spe1-HF (NEB #R3133) restriction enzymes to generate the eGATA3 expression plasmid. Stable transfected T47D cell lines were generated using Lipofectamine 2000 (Invitrogen, 11668030) according to the manufacturer’s instructions. Stable transduced cell lines were generated by lentiviral transduction using the psPAX2 (Addgene plasmid #12260; RRID:Addgene_12260) (500 ng) and pMD2.G (Addgene plasmid #12259; RRID:Addgene_12259) (500 ng) and eGATA3 plasmid (750 ng) plasmids. Stable cell populations were selected with puromycin (2 µg/mL) and subsequently maintained in a medium containing 1 µg/mL puromycin.

### Western blotting

Cells were washed with phosphate-buffered saline (PBS Gibco 70011-044), scraped from 30 mm culture dishes, and lysed directly in 2X Laemmli sample buffer. Lysates were heated at 95°C for 5 min and clarified by centrifugation at 13,400 rpm for 1 min. Proteins were separated by SDS-PAGE using 4% stacking and 10% resolving gels and transferred onto polyvinylidene fluoride (PVDF) membranes (Millipore, IPVH00010) in transfer buffer (25 mM Tris, 192 mM glycine, and 20% methanol) at 20 V for 1 h. Membranes were blocked with 5% skimmed milk (Santa Cruz Biotechnology, sc-2324) in TBS-T for 1 h at room temperature and incubated overnight at 4°C with rabbit anti-GATA3 (Cell Signaling Technology D13C9; 1:2000) and mouse anti-beta actin (Santa Cruz Biotechnology, sc-47778; 1:5000) primary antibodies. After washing with TBS-T, membranes were incubated with horseradish peroxidase-conjugated anti-rabbit (Invitrogen, 32460; 1:10,000) or anti-mouse (Invitrogen, 32430; 1:10,000) secondary antibodies for 1 h at room temperature. Immunoreactive bands were detected using Clarity Western ECL substrate (Bio-Rad, 1705060), imaged using an ImageQuant LAS 4000 imaging system, and processed using ImageJ software.

### Mammosphere formation assay

Mammosphere culture was performed as previously described^75^. T47D cell lines were dissociated into single-cell suspensions by trypsinization using 0.25% of trypsin-EDTA. In ultra-low attachment 96-well plates (Corning), 1-3 cells were seeded per well. Cells were maintained in serum-free DMEM F-12 supplemented with 1% glutamine, 1% Penstrep, 2% B27, 20 ng/ml EGF, and 20 ng/ml FGFb. Mammosphere cultures were then incubated at 37 degrees and 5% CO₂ for a period of one week. Three independent biological replicates were performed, with 12 wells per condition per replicate. Wells were inspected by phase contrast microscopy, and images were acquired by a Nikon ECLIPSE T2000 inverted microscope. Mammospheres are defined by a size of 50um or above. Sphere formation efficiency (SFE) is calculated as the proportion of seeded wells that contain one mammosphere. Diameters were quantified by the NIS Elements software.

### Spheroid Assay

A total of 5,000 cells were seeded per well in 96-well ultra-low attachment plates (Corning). Three independent biological replicates were performed, with 7 wells per condition per replicate. Spheroid formation was monitored at 24 h and 48 h. Brightfield images were acquired using a Nikon ECLIPSE T2000 inverted microscope equipped with the 10X objective. Spheroid morphology was quantified by calculating circularity using NIS-Elements software (Nikon).

### RNA sequencing

RNA from T47D lines was extracted using Trizol, quantified by Qubit fluorometry, and RNA integrity was assessed by Bioanalyzer (RIN > 8). The NEBNext Poly(A) mRNA Magnetic Isolation Module (Cat: E7490L) was used to capture mRNA, and libraries were prepared using the NEBNext Ultra II Directional RNA Library Prep kit (Cat: E7765L). Six independent biological replicates were performed per condition, and the libraries were sequenced on the NovaSeq Platform with a target of approximately 50 million reads per sample. For MCF7, library preparation and sequencing were carried out by the Genomics Core Facility (CRUK-CI). RNA libraries were prepared using the TruSeq stranded mRNA library prep kit (Illumina), and samples were sequenced on a HiSeq 4000 to approximately 30 million reads per sample.

Published RNA-seq data from GATA3 ZnF2-mutant cells were obtained from the Gene Expression Omnibus (GEO; GSE99479) and reanalyzed in this study. Fastq reads were processed using the nf-core RNA-Seq pipeline (version v3.14.0), a standardized and automated pipeline designed for comprehensive RNA-Seq data analysis^76^. RNA-seq reads quantified using Salmon were imported into DESeq2 using tximport. Differential gene expression analysis was carried out through DESeq2 (version 1.46.0)^77^. DEGs were defined at thresholds of |log_2_FC| > 1.5 and FDR < 0.01.

### CUT&RUN

Cleavage Under Targets and Release Using Nuclease (CUT&RUN) was performed following the protocol described by Derek Janssens and Steven Henikoff (version 3)^78^. Briefly, 200,000 cells per reaction were harvested and immobilized on concanavalin A magnetic beads (BioMag®Plus Concanavalin A, BP531-3). Cells were gently permeabilized to maintain nuclear integrity while allowing antibody access. Samples were incubated with a primary antibody against GATA3 (Cell Signaling Technology, GATA3-D13C9) along with an IgG control antibody included in the Cell Signaling Technology CUT&RUN kit (66362S). Following primary antibody binding, the pA-MNase enzyme (Cell Signaling Technology kit, 57813S) was added to target antibody-bound chromatin. After appropriate washing steps to remove unbound enzyme, controlled digestion was initiated by calcium addition, enabling precise cleavage of DNA adjacent to protein-binding sites. The reaction was terminated using stop buffer, and released DNA fragments were collected from the supernatant. A defined amount of spike-in DNA of yeast origin (100 pg per reaction; Cell Signaling Technology kit, 36598S) was included to enable normalization across samples. DNA was purified using the phenol-chloroform extraction method and processed for downstream applications, including library preparation and sequencing with a target of around 18M reads per sample sequenced on Novaseq platform.

### CUT&RUN data analysis

CUT&RUN sequencing data were processed using the nf-core/cutandrun pipeline (version 3.2.2) implemented in Nextflow (version 24.10.0)^76^. Reads were aligned to the human reference genome (hg38) with spike-in normalization using the Saccharomyces cerevisiae R64-1-1 reference genome. Duplicate reads were removed, and reads with a mapping quality score <20 were excluded. Peak calling was performed using both MACS2 (q-value < 0.05) and SEACR (stringent mode). Genome browser tracks were visualized using the Integrative Genomics Viewer (IGV)^79^.

### FOXA1 ChIP-seq in MCF7

MCF7 cells (control and eGATA3 mutant)^37^ were cross-linked for 10 minutes using 1% formaldehyde (Thermo Scientific, ref. 28908) and quenched with 0.1M glycine. All the following lysis buffers were supplemented with a protease inhibitor cocktail. Pellets were resuspended in Lysis Buffer-1 (50 mM Hepes–KOH, pH 7.5, 140 mM NaCl, 1 mM EDTA, 10% Glycerol, 0.5% NP-40/Igepal CA-630, 0.25% Triton X-100, and rotated for 10 minutes at 4°C. Cells were then pelleted and resuspended in Lysis buffer-2 (10 mM Tris–HCl, pH 8.0, 200 mM NaCl, 1 mM EDTA, 0.5 mM EGTA) and incubated for 5 minutes at 4°C with rotation. Cells were then pelleted and resuspended in 300 µl Lysis buffer-3 per 15 cm² plate (10 mM Tris–HCl, pH 8, 100 mM NaCl, 1 mM EDTA, 0.5 mM EGTA, 0.1% Sodium deoxycholate) and sonicated using the Bioruptor® Plus sonicator (Diagenode, Liege, Belgium) for 15 cycles (30 seconds on, 30 seconds off). After sonication, 1% Triton X was added to the samples, after which they were centrifuged at maximum speed for 10 minutes at 4°C and a small aliquot of supernatant was kept as input for ChIP. The rest of the supernatant was added to the antibody-bead conjugate after adequate washing with Bovine Serum Albumin made in PBS and resuspended in Lysis buffer-3. For each ChIP, 10 µg of the appropriate antibody (ab5089) was added to 100 µl of Dynabeads® Protein A or G for the rabbit or goat antibodies, respectively. Each antibody and beads were incubated together overnight at 4°C. The beads after incubation were washed six times with a modified RIPA buffer (150 mM NaCl, 10 mM Tris, pH 7.2, 0.1% SDS, 1% Triton X-100, 1% sodium deoxycholate), followed by 10 mM Tris-1 mM EDTA buffer pH 7.4. Both ChIP samples and inputs were then de-crosslinked in 200 µl elution buffer (1% SDS, 0.1 M NaHCO₃) overnight at 65°C. After reverse crosslinking, DNA was isolated and purified using the phenol-chloroform-isoamyl DNA extraction method. ChIP-seq and the input libraries were prepared using the ThruPlex® DNA-seq kit and sequenced on HiSeq 2500. ChIP-seq was performed in biological triplicates, using cells from independent passages. The GATA3 and ERα ChIP-seq datasets performed in the same cell lines were obtained from the previous study (GSE153255)^37^.

### ChIP-seq data analysis

Raw sequencing data were processed using the nf-core/chipseq pipeline^76^, including read quality control, adapter trimming, alignment to the reference genome (hg38), and peak calling.

Differential binding was assessed using DiffBind (v3.16.0), with DESeq2 as the underlying statistical framework. Biological replicates were included as independent samples within each condition and analyzed jointly. Peaks were quantified using dba.count with a summit size of 250 bp, and differential binding was assessed using a condition-based contrast. Default library-size normalization was applied, followed by differential analysis using DESeq2. The complete set of differential binding results was exported for downstream analysis. Gained and lost GATA3-binding sites were subsequently defined using the FDR threshold of 0.05.

### Single-nuclei mRNA and ATAC sequencing

Single-cell chromatin accessibility and gene expression profiling were performed using the 10X Genomics Chromium Next GEM Single Cell Multiome ATAC + Gene Expression kit, according to the manufacturer’s protocols. Briefly, nuclei were isolated from cells using buffer compositions and procedures as described in the kit guidelines, ensuring preservation of chromatin structure and RNA integrity. A total of 16,000 nuclei per reaction were used as input for the assay. Isolated nuclei were loaded onto the Chromium controller to generate Gel Bead-In-Emulsions, enabling parallel barcoding of individual nuclei for both ATAC (chromatin accessibility) and gene expression (GEX) profiling. Following transposition and reverse transcription within GEMs, libraries were constructed according to the manufacturer’s instructions. The resulting ATAC and gene expression libraries were sequenced and processed to obtain genome-wide chromatin accessibility profiles and transcriptomic data at single-cell resolution.

### Single-nuclei multiome data analysis

Raw sequencing data were processed using Cell Ranger ARC (10x Genomics) to generate paired gene expression and chromatin accessibility count matrices. Single-nucleus RNA-seq data were analyzed using Scanpy (version 1.11.5)^80^. Following quality-control filtering, gene expression data were normalized, log-transformed, and subjected to principal component analysis, neighborhood graph construction, and Uniform Manifold Approximation and Projection (UMAP) for dimensionality reduction and visualization. Differential gene expression analysis and gene signature scoring were performed using Scanpy. Single-nucleus ATAC-seq data were analyzed using SnapATAC2 (version 2.9.0)^81^. Following quality-control filtering; dimensionality reduction, clustering, and differential chromatin accessibility analyses were performed using the standard snapATAC2 workflow. Differentially accessible regions were subsequently used for transcription factor motif enrichment analysis using HOMER. Trajectory inference was performed using partition-based graph abstraction (PAGA) following diffusion pseudotime calculation. G2/M-phase cells were excluded to minimize the influence of cell-cycle-associated transcriptional variation^82^. Wild-type luminal cells were defined as the root population for trajectory inference, and lineage-associated gene signature scores were projected along the inferred pseudotime trajectory.

### Structure prediction with AlphaFold3

Structures for the wild-type GATA3 and eGATA3 dimers in complex with the GATA3 DNA motif were obtained using the AlphaFold3 (AF3) webserver (https://alphafoldserver.com/)^49^. For both the complexes, we obtained AF3 predictions using five random seeds, with five structures predicted for each seed, resulting in a total of 25 predicted structures. The best predicted structure was selected based on the ranking score provided by AF3. We identified the confident regions from the best predicted structure using the open-source AF-Pipeline, with PAE and pLDDT thresholds of 12 and 70, respectively (https://github.com/isblab/af_pipeline).

## Acknowledgement

We acknowledge the funding support from the Department of Atomic Energy, Government of India, under Project Identification No. RTI 4006 and intramural funds from NCBS-TIFR. RS acknowledges support from the DBT/Wellcome Trust India Alliance Fellowship [grant number IA/I/20/1/504928]. We thank Hisham Mohammed, Jyothi S. Prabhu, Shivaprasad PV, Anurag Kumar Singh, Deepanshu Soota and members of RS lab for their feedback and suggestions on this manuscript.

## Author contributions

NV and RS conceived and designed the study. NV performed most of the experiments, data curation and analysis, interpreted the results and prepared figures. MR contributed to the CUT&RUN and single-nuclei multi-omics experiments, data analysis and interpretation. KM and SV contributed to the in silico protein structure prediction analysis. SJ, IC and JSC contributed to the ChIP-seq and RNA sequencing of the MCF7 cell line. DN contributed to the experimental design, interpreted the results and provided feedback. NV wrote the manuscript with input from all authors. RS supervised the study. All authors read and approved the final manuscript.

