## Supplementary Figures for "GATA3-extension mutants rewire lineage identity in luminal breast cancer"

**Supplementary Figure S1**

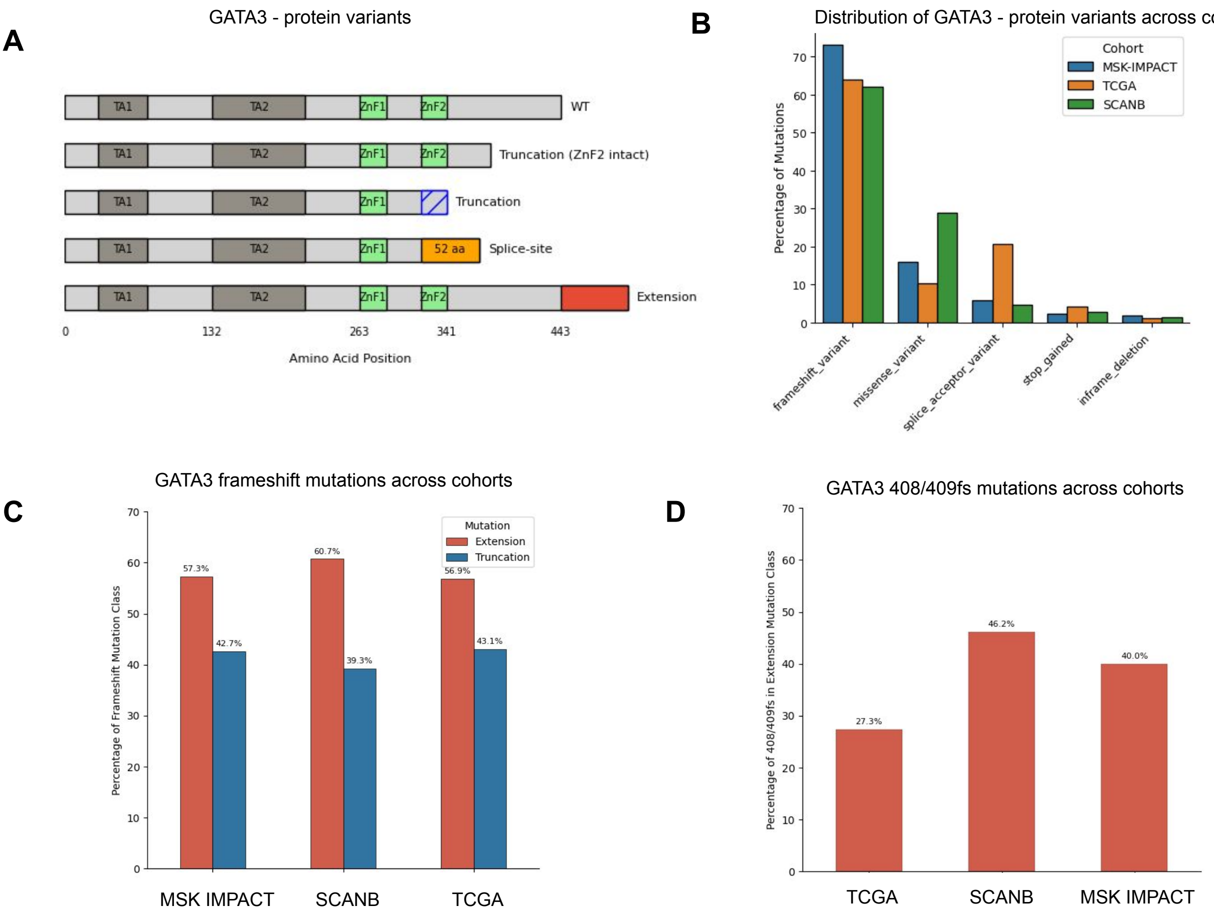


**Figure S1. Distribution of GATA3 mutations across breast cancer cohorts**

**(A)** Schematic representation of the major GATA3 protein variants generated by frameshift mutations. **(B)** Distribution of protein-altering GATA3 mutations across the MSK-IMPACT, TCGA, and SCAN-B breast cancer cohorts. Mutation classes are grouped according to their annotated variant consequence. **(C)** Proportion of frameshift mutations classified as extension or truncation mutations in the MSK-IMPACT, TCGA-BRCA, and SCAN-B cohorts. **(D)** Frequency of recurrent GATA3 p.P408/409fs mutations among extension mutations in the MSK-IMPACT, TCGA-BRCA, and SCAN-B cohorts.

### Supplementary Figure S2

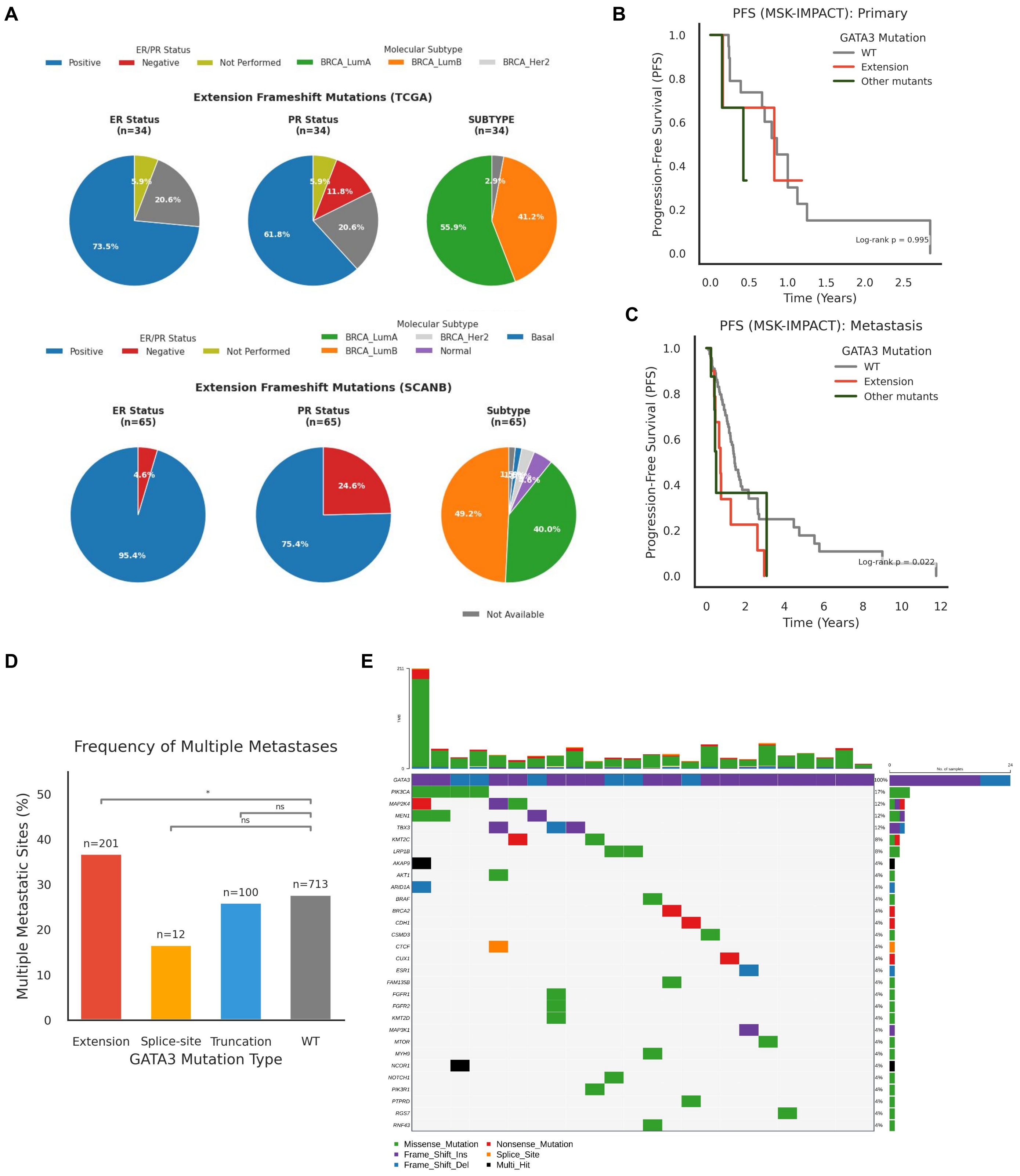

**Figure S2. Clinical and molecular associations of GATA3 mutants in breast cancer**

**(A)** Distribution of receptor status and subtypes among eGATA3-mutant tumors in the TCGA-BRCA and SCAN-B cohorts. Percentages indicate the proportion of tumors assigned to each category. **(B-C)** Kaplan–Meier analyses of progression-free survival (PFS) in primary (B) and metastatic (C) breast tumors from the MSK-IMPACT cohort stratified by GATA3 mutation status. Statistical significance was assessed using a pairwise log-rank test comparing eGATA3-mutant and wild-type tumors. Other mutants comprise truncating and splice-site GATA3 mutants and are shown for comparison only. **(D)** Percentage of extension, truncation, and splice-site GATA3 mutations with multiple metastases in the MSK-IMPACT cohort with respect to the WT tumors. Statistical significance was assessed using Fisher's exact test. **(E)** Recurrently co-mutated cancer driver genes identified in patients harbouring extension GATA3 mutations. Bars indicate the number of patients carrying alterations in each corresponding driver gene. 'Multi_Hit' denotes genes harbouring two or more mutations within the same tumor sample. Asterisks denote statistical significance: $P < 0.05$ (*); ns, not significant.

#### Supplementary Figure S3

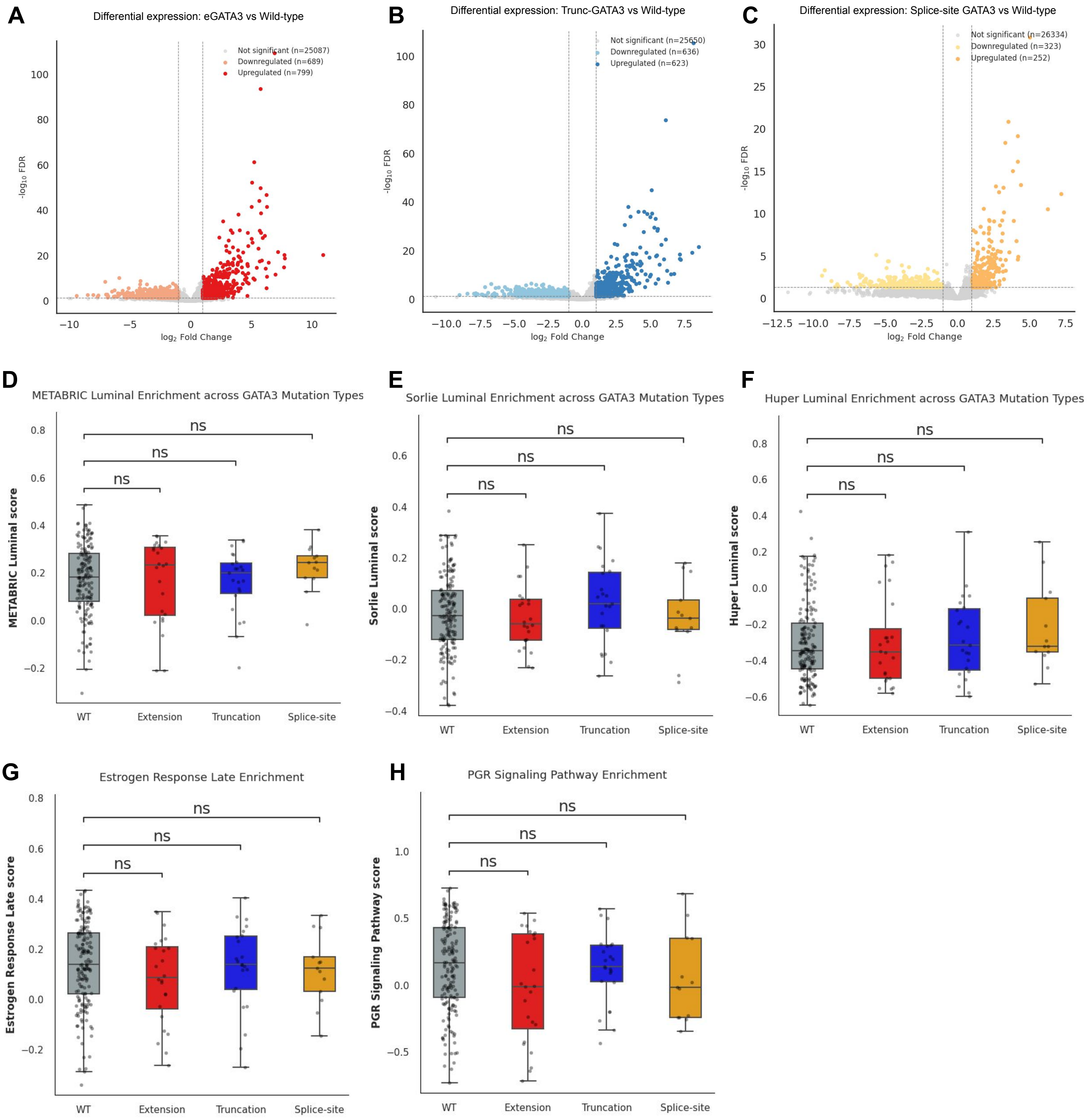


**Figure S3. Mutation class-specific transcriptional alterations with respect to WT tumors (A–C)** Differential gene expression in extension (A), truncating (B), and splice-site (C) mutants relative to GATA3 wild-type tumors in the TCGA cohort. Significantly upregulated and downregulated genes are highlighted. **(D-I)** GSVA scores for the METABRIC luminal (D), Sorlie luminal (E), Huper luminal (F), Estrogen Response Late (G), and Progesterone Receptor (PGR) Signalling (H) signatures across GATA3 mutants in TCGA breast tumors. In boxplots, boxes denote the interquartile range (Q1–Q3), the central line indicates the median, and whiskers extend to 1.5× the interquartile range. Data points beyond the whiskers are displayed as outliers. Statistical significance was assessed using the Mann-Whitney U test (two-sided). GSVA scores of FDR < 0.05 were considered significant. Asterisks denote statistical significance: $P < 0.05$ (*), $P < 0.01$ (**), and $P < 0.001$ (***); ns, not significant.

#### Supplementary Figure S4

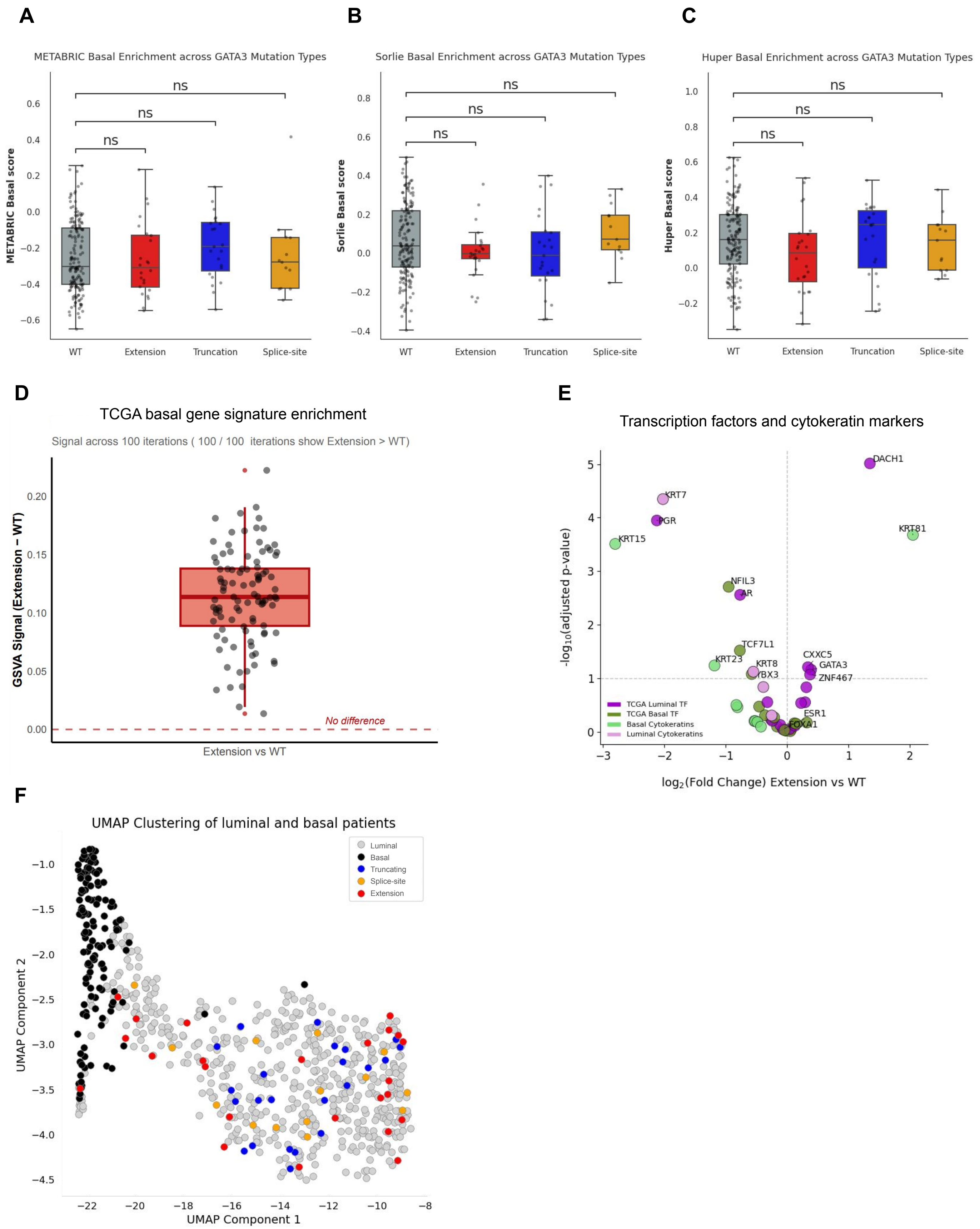

**Figure S4. Additional characterization of the transcriptional phenotype associated with eGATA3 mutations.**

**(A-C)** GSVA enrichment scores for the METABRIC basal (A), Sorlie basal (B), and Huper basal (C) gene signatures across GATA3 mutation classes. **(D)** Bootstrap analysis (100 iterations) demonstrating robust enrichment of the TCGA basal gene signature in eGATA3-mutant tumors compared with wild-type tumors. Each point represents the GSVA score obtained from one bootstrap iteration. **(E)** Differential expression analysis of cytokeratin genes and TCGA-derived lineage-associated TFs in eGATA3-mutant tumors with respect to WT tumors. Dashed lines indicate the differential expression thresholds used for significance. **(F)** UMAP projection of TCGA breast tumors using the TCGA basal gene signature, showing the distribution of wild-type and GATA3-mutant tumors. Boxes represent the interquartile range (IQR), center lines indicate the median, whiskers extend to 1.5 × IQR, and points represent individual tumors. Statistical significance was assessed using the Mann–Whitney U test with Benjamini–Hochberg correction; $P < 0.05$, P < 0.01, $P < 0.001$; ns, not significant.

#### Supplementary Figure S5

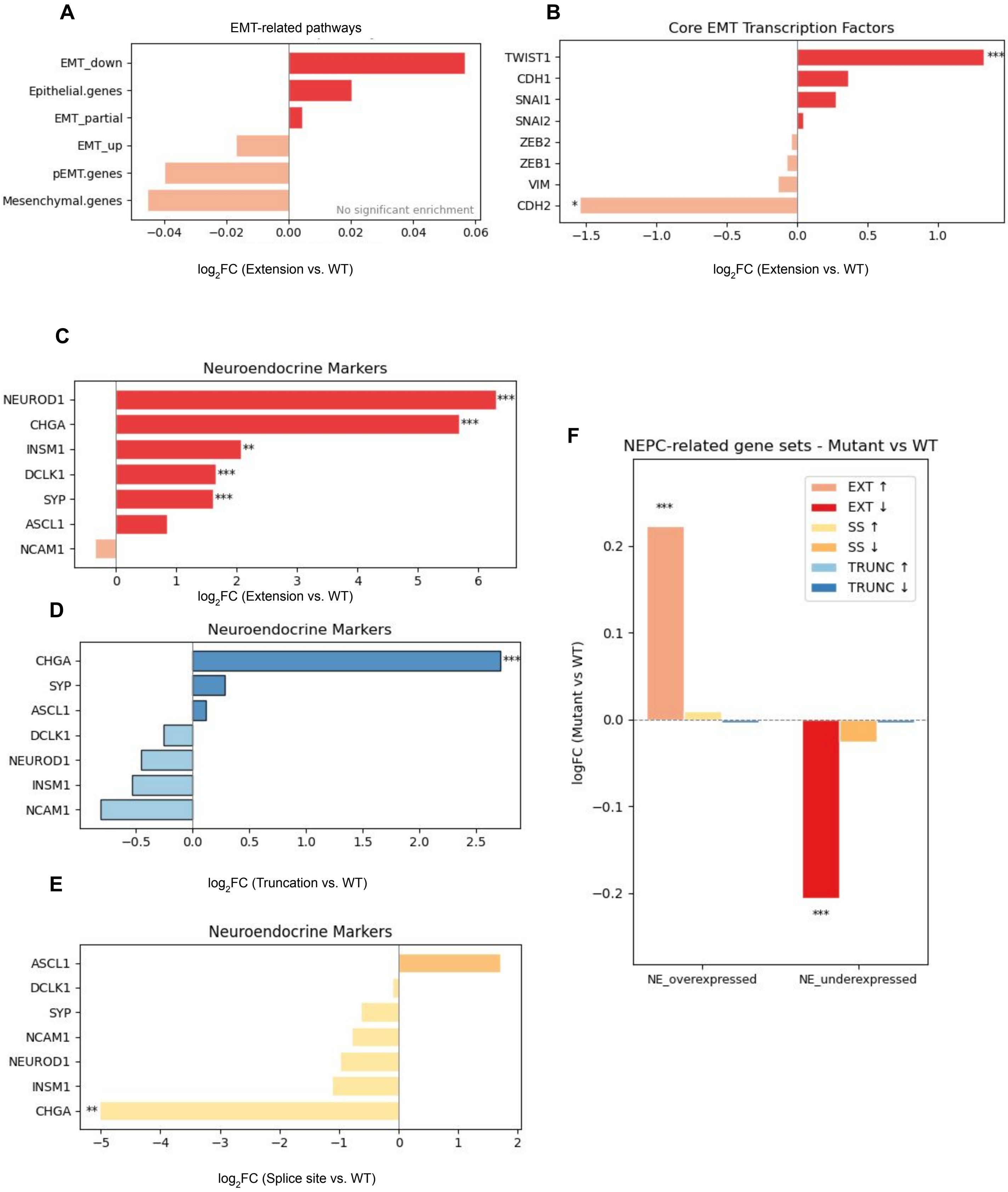

**Figure S5. eGATA3 promotes neuroendocrine-associated transcription without activating a canonical EMT program.**

**(A)** Differential enrichment analysis of epithelial–mesenchymal transition (EMT)-related gene signatures in eGATA3-mutant tumors relative to wild-type tumors. **(B)** Differential expression of core epithelial-mesenchymal transition (EMT) transcription factors in eGATA3-mutant tumors relative to WT tumors. Only TWIST1 was significantly upregulated, whereas other canonical EMT regulators remained largely unchanged. **(C–E)** Differential expression of neuroendocrine marker genes in extension (C), truncating (D), and splice-site (E) GATA3 mutant tumors relative to wild-type tumors. eGATA3 mutants display the strongest induction of neuroendocrine-associated markers. **(F)** Differential enrichment of GSVA scores of curated neuroendocrine prostate cancer (NEPC)-associated gene signatures across GATA3 mutation classes. Extension mutants exhibit enrichment of neuroendocrine-overexpressed genes together with depletion of neuroendocrine-underexpressed genes, supporting activation of a neuroendocrine-associated transcriptional program. Genes with FDR < 0.05 were considered differentially expressed. Asterisks denote statistical significance: $P < 0.05$ (*), $P < 0.01$ (**), and $P < 0.001$ (***); ns, not significant.

#### Supplementary Figure S6

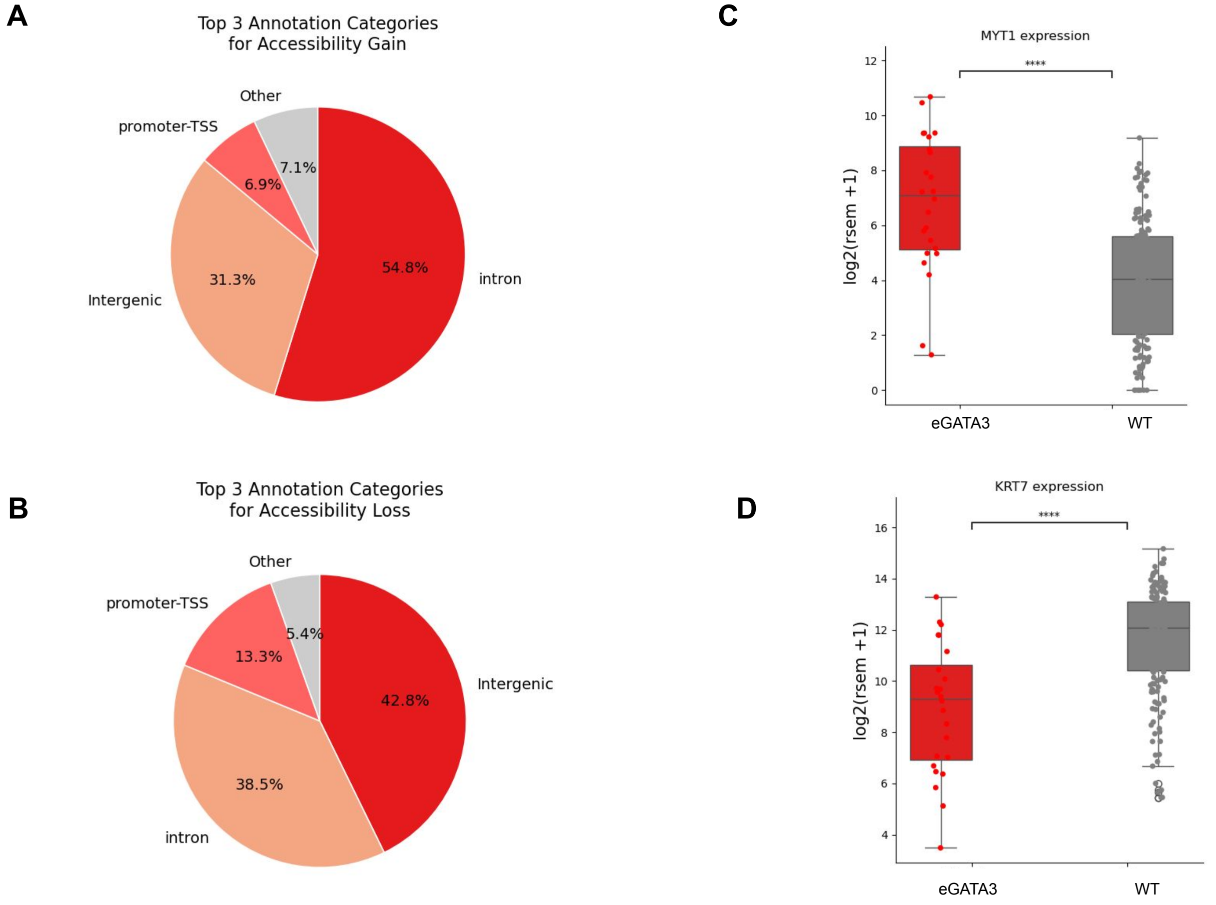


**Figure S6. Genomic annotation and representative changes associated with eGATA3 chromatin remodeling.**

**(A)** Genomic annotation of regions exhibiting increased chromatin accessibility in eGATA3-mutant tumors relative to GATA3 wild-type tumors. **(B)** Genomic annotation of regions exhibiting decreased chromatin accessibility. **(C)** Expression of **MYT1** in eGATA3-mutant and GATA3 wild-type tumors. **(D)** Expression of **KRT7** in eGATA3-mutant and GATA3 wild-type tumors.

#### Supplementary Figure S7

A

TOP MOTIFS ENRICHED IN GAINED

| MOTIF | Best Match | p-value | % targets |
|---|---|---|---|
| 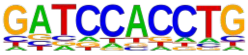 | E2A | 1e-13 | 51.19% |
| 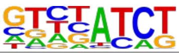 | GATA3 | 1e-15 | 38.58% |
| 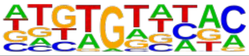 | FOXO3 | 1e-31 | 24.35% |

C

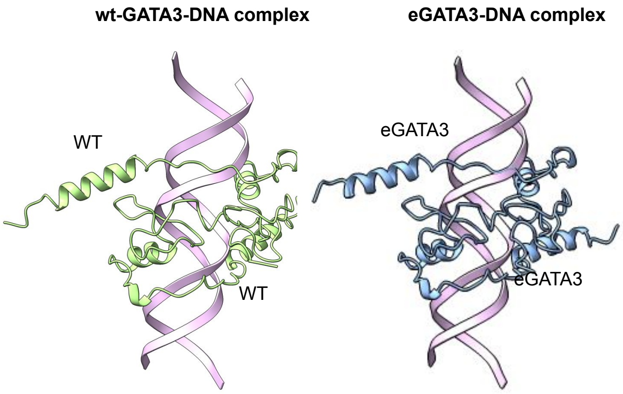


B

TOP MOTIFS ENRICHED IN LOSS

| MOTIF | Best Match | p-value | % targets |
|---|---|---|---|
| 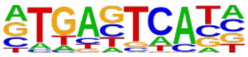 | FRA1 | 1e-84 | 37.83% |
| 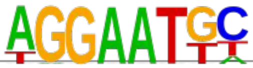 | TEAD | 1e-31 | 26.55% |
| 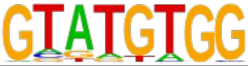 | RUNX2 | 1e-12 | 15.59% |

D

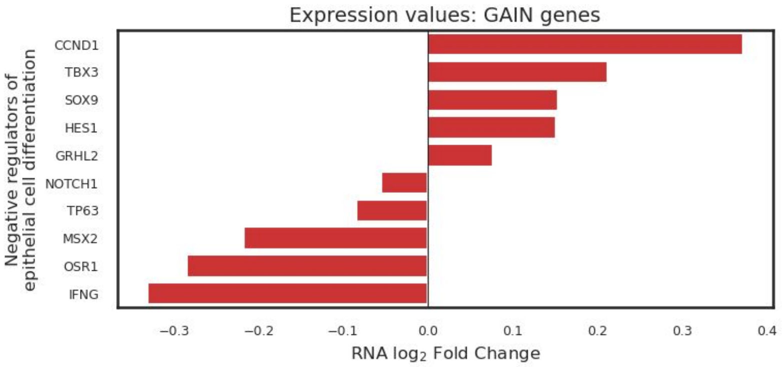


E

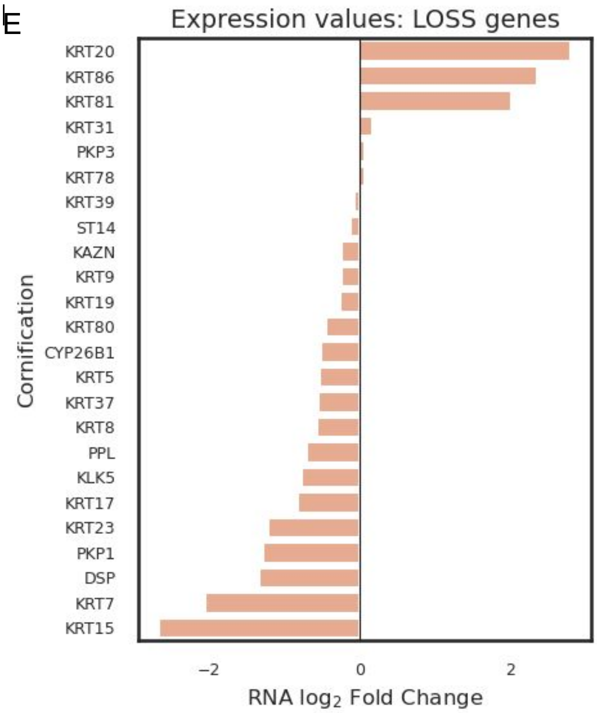


**Figure S7: Motif and structural analyses support eGATA3-associated chromatin remodelling.**

**(A)** Top enriched transcription factor motifs identified in regions exhibiting increased chromatin accessibility. **(B)** Top enriched transcription factor motifs identified in regions exhibiting decreased chromatin accessibility. **(C)** AlphaFold predicted structures for the wild-type and eGATA3–DNA complexes. **(D)** Differential expression of representative genes from the top enriched Gene Ontology pathway associated with regions exhibiting increased chromatin accessibility. **(E)** Differential expression of representative genes from the top enriched Gene Ontology pathway associated with regions exhibiting decreased chromatin accessibility.

### Supplementary Figure S8

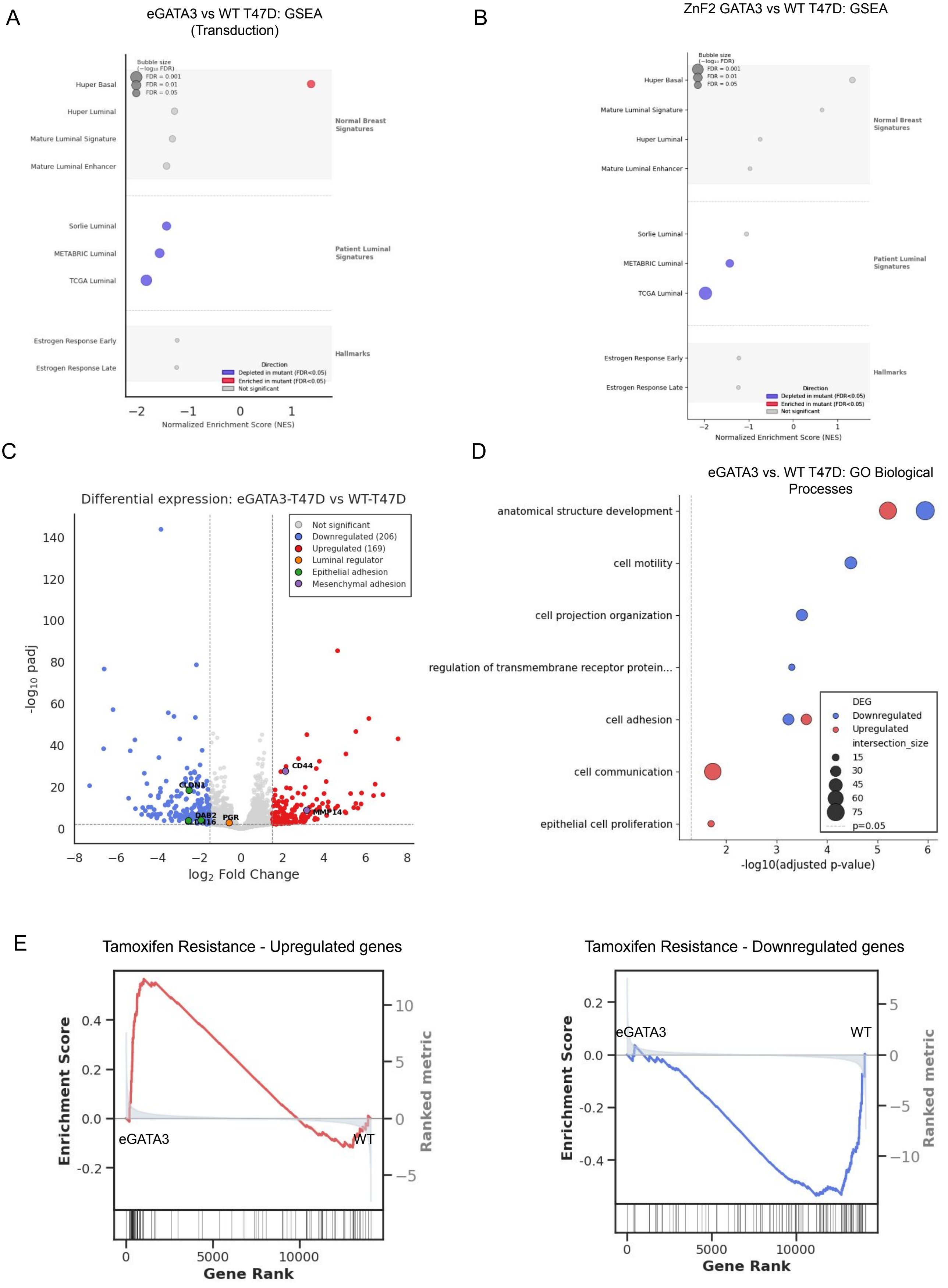

**Figure S8: Additional transcriptomic characterization of eGATA3-expressing T47D cells.**

**(A):** Gene set enrichment analysis (GSEA) of RNA sequencing data from the independently generated eGATA3-transduced T47D cell line relative to wild-type cells **(B)** Gene set enrichment analysis of published RNA-sequencing data from ZnF2-truncating GATA3-expressing T47D cells relative to wild-type cells. **(C)** Volcano plot showing differentially expressed genes in eGATA3-expressing T47D cells relative to wild-type cells. Selected genes associated with luminal identity, epithelial cell adhesion, and mesenchymal cell adhesion are highlighted. **(D)** Gene Ontology enrichment analysis of differentially expressed genes in eGATA3-expressing T47D cells **(E)** Gene set enrichment analysis demonstrating enrichment of genes upregulated in tamoxifen-resistant T47D cells and depletion of genes downregulated in tamoxifen-resistant T47D cells.

#### Supplementary Figure S9

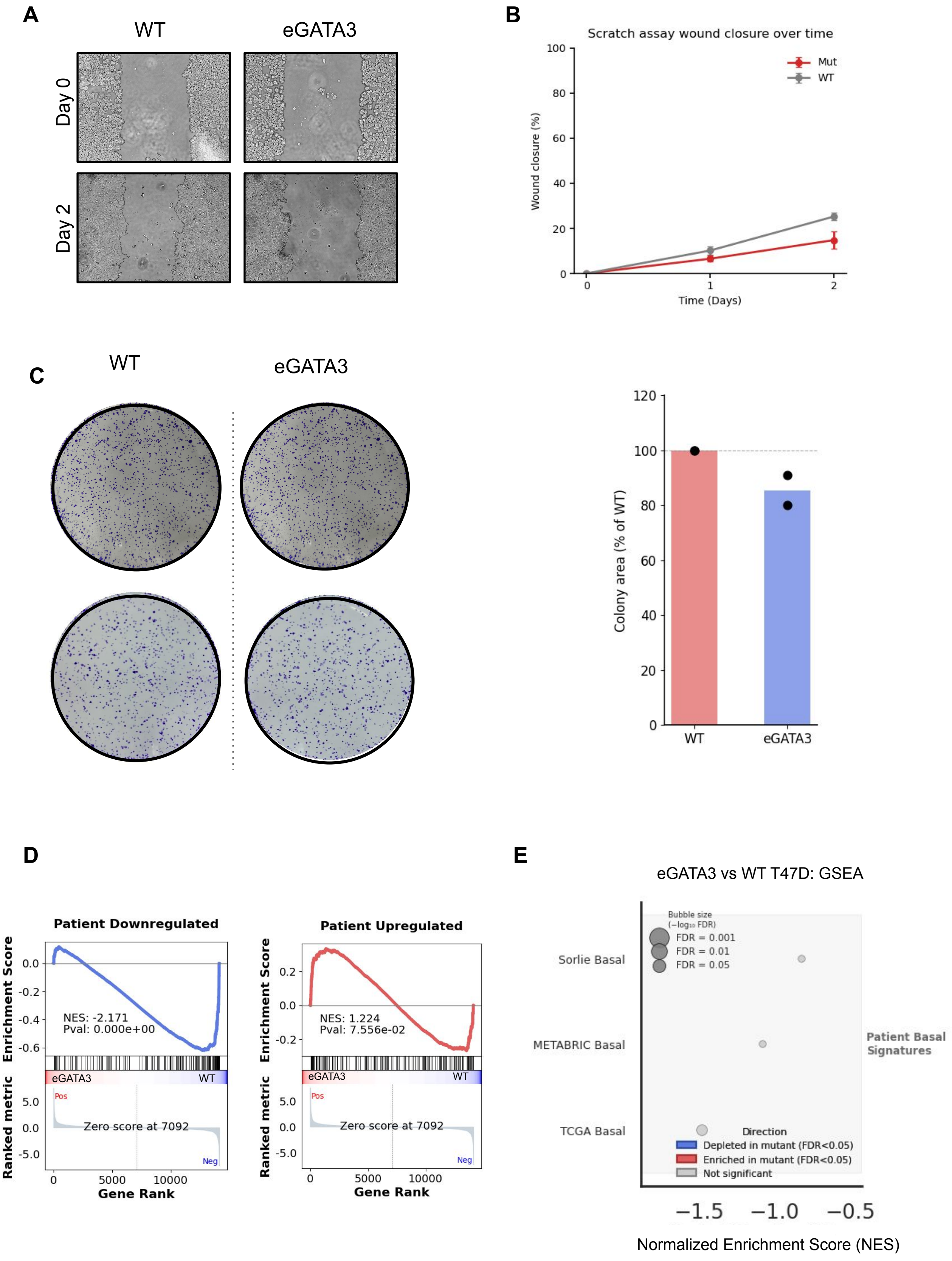

**Figure S9: Functional characterization and comparison of eGATA3 transcriptional programs with patient tumors.**

**(A)** Representative brightfield images from wound-healing assays performed in wild-type and eGATA 3-expressing T47D cells at the indicated time points. **(B)** Quantification of wound closure between wild-type and eGATA3-expressing cells. **(C)** Representative crystal violet-stained colony formation assays  and quantification of colony area. **(D)** Gene set enrichment analysis comparing genes that are differentially expressed in eGATA3-mutant patient tumors in the eGATA3 T47D transcriptome. **(E)** Gene set enrichment analysis of RNA-sequencing data from eGATA3-transfected T47D cells relative to wild-type cells for the tumor derived basal signatures.

#### Supplementary Figure S10

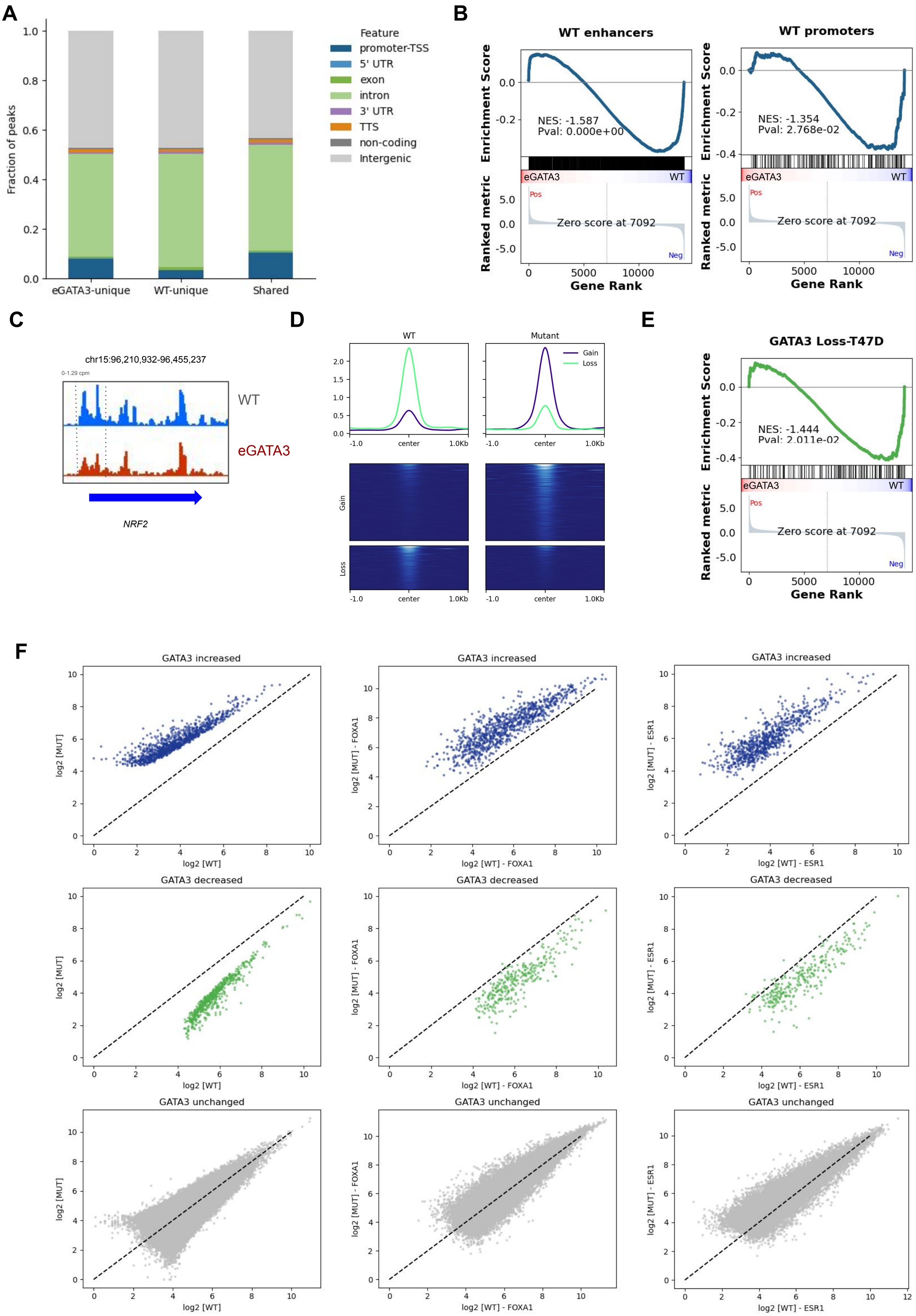

**Figure S10: Redistribution of GATA3 occupancy following eGATA3 over-expression in breast cancer cells.**

**(A)** Genomic annotation of wild-type-unique, eGATA3-unique, and shared GATA3 binding sites identified by CUT&RUN, showing the distribution of peaks across promoters, untranslated regions (UTRs), exons, introns, transcription termination sites (TTSs), and intergenic regions. **(B)** Gene set enrichment analysis demonstrating downregulation of genes associated with wild-type-unique promoter and enhancer GATA3 binding sites in eGATA3-expressing T47D cells. **(C)** Representative genome browser tracks illustrating loss of GATA3 occupancy at the NRF2 locus in eGATA3-expressing T47D cells relative to wild-type cells. **(D)** Average GATA3 ChIP signal in MCF7 at gain and loss GATA3 binding sites. **(E)** Gene set enrichment analysis demonstrating depletion of genes associated with regions losing GATA3 occupancy in MCF7 in the eGATA3 T47D transcriptome. **(F)** Average signal from FOXA1 and ER ChIP-seq across regions exhibiting increased, decreased, or unchanged GATA3 occupancy. Regions gaining GATA3 occupancy show increased FOXA1 and ER signal, whereas regions losing GATA3 occupancy exhibit reduced FOXA1 and ER signal.

### Supplementary Figure S11

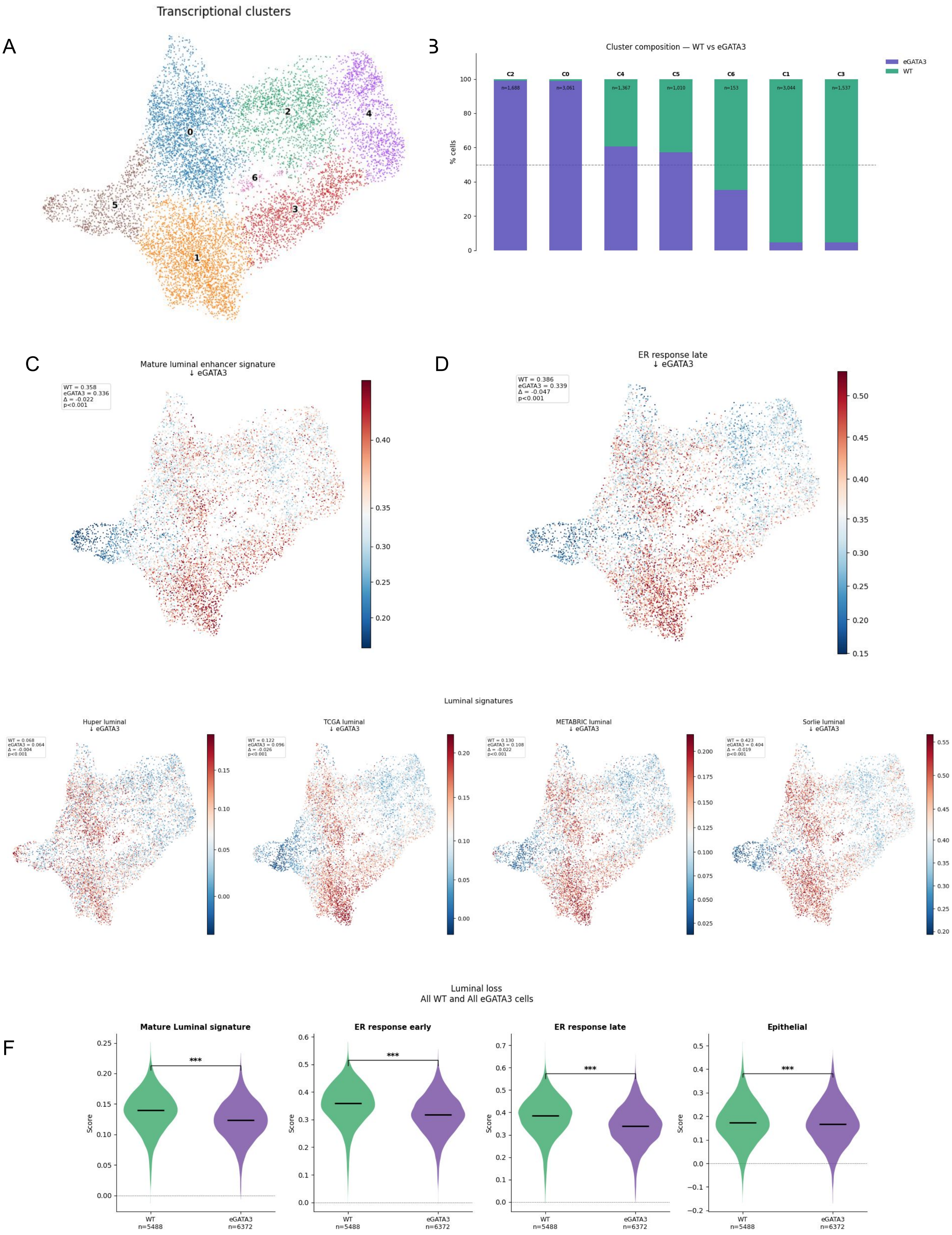

**Figure S11: Single-nucleus transcriptomic characterization of eGATA3-expressing T47D cells.**

**(A)** UMAP of snRNA-seq profiles showing transcriptionally defined clusters identified by unsupervised clustering. **(B)** Relative contribution of wild-type and eGATA3-expressing cells to each transcriptional cluster. **(C)** Projection of the mature luminal enhancer signature onto the snRNA-seq UMAP. **(D)** Projection of the estrogen-response late signature onto the snRNA-seq UMAP. **(E)** Projection of additional lineage-associated gene signatures, including Huper luminal, TCGA luminal, METABRIC luminal, and Sørlie luminal signatures. **(F)** Violin plots comparing single-cell signature scores for mature luminal, estrogen-response early, estrogen-response late and epithelial gene signatures between wild-type and eGATA3-expressing cells. Statistical significance was assessed using the two-sided Mann–Whitney U test; $P < 0.05$, *$P < 0.01$, $P < 0.001$, ns, not significant.

#### Supplementary Figure S12

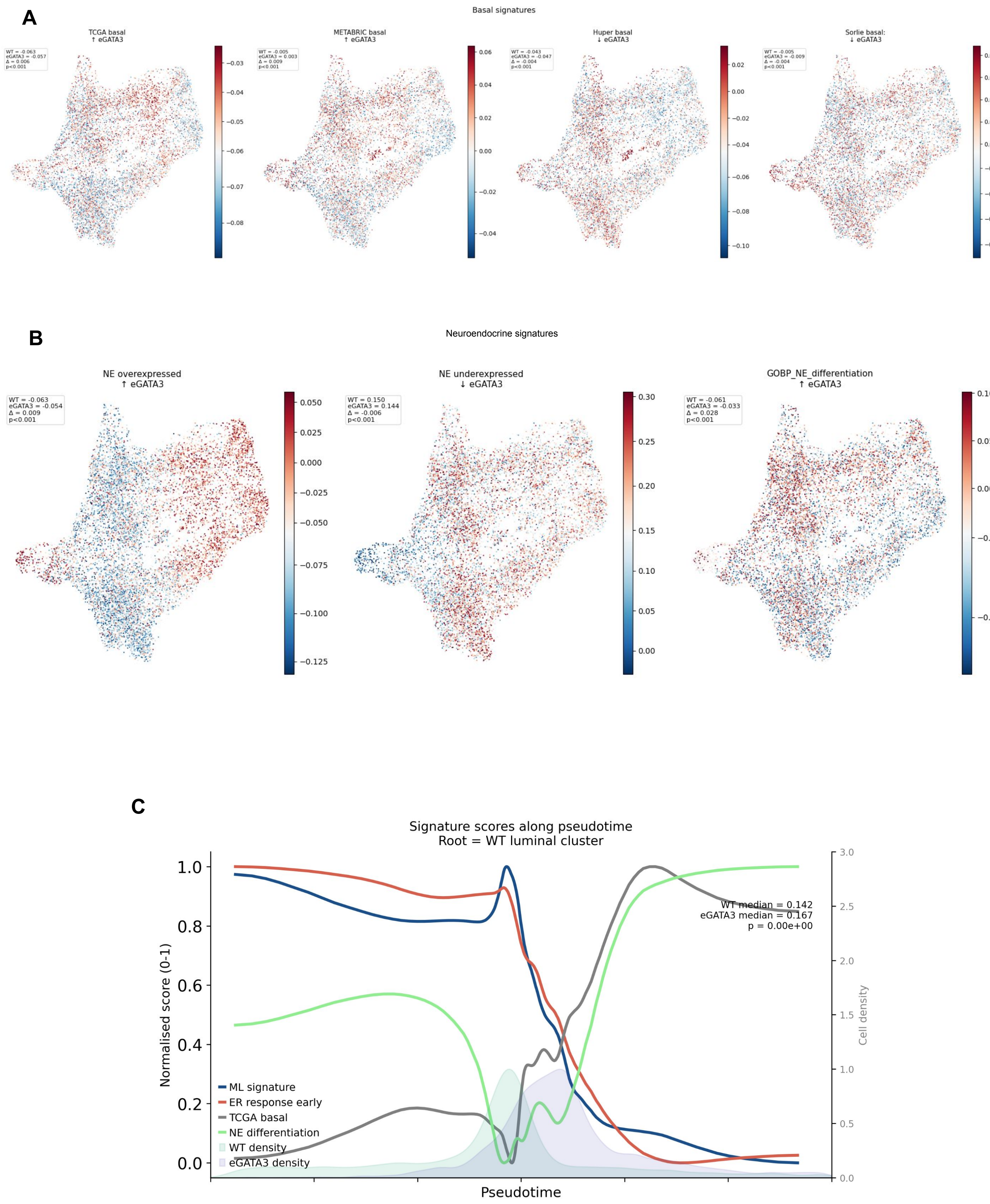

**Figure S12: Single-cell analyses supporting eGATA3-associated lineage reprogramming.**

(A) Projection of additional basal-associated gene signatures onto the snRNA-seq UMAP, including TCGA basal, METABRIC basal, Huper basal, and Sørlie basal signatures. (B) Projection of neuroendocrine (NE)-associated gene signatures onto the snRNA-seq UMAP, including neuroendocrine overexpressed, neuroendocrine underexpressed, and GO: neuroendocrine differentiation gene signatures. (C) Integrated Diffusion Pseudotime analysis showing progressive changes in mature luminal, estrogen-response, TCGA basal, and neuroendocrine-associated transcriptional programs along the inferred transcriptional trajectory. Statistical significance was assessed using the two-sided Mann–Whitney U test; $P < 0.05$, *$P < 0.01$, $P < 0.001$, ns, not significant.
